# Scalable enumeration and sampling of minimal metabolic pathways for organisms and communities

**DOI:** 10.1101/2024.10.09.617357

**Authors:** Ove Øyås, Axel Theorell, Jörg Stelling

## Abstract

Many interactions in microbial consortia or tissues of multicellular organisms rely on networks of metabolite exchanges. To predict community function and composition beyond statistical correlations, one can use genome-scale metabolic models. However, comprehensive model analysis via metabolic pathways is a major challenge because pathway counts grow combinatorially with model size. Here, we define minimal pathways that yield compact representations of metabolic network capabilities. They generalize existing pathway concepts by allowing inhomogeneous constraints and targeted analysis of subnetworks, and we show how to enumerate and sample them efficiently via iterative minimization and pathway graphs. This enables applications such as assessing quantitative gene essentiality in the central metabolism of *Escherichia coli*, predicting metabolite exchanges associated with homeostasis and health in a host-microbe model of the human gut, and designing butyrate-producing microbial communities. Minimal pathways enable scalable analysis of metabolic subnetworks such as metabolite exchanges in uni- and multicellular systems.

## Introduction

Many functions of microbial communities and of tissues in multicellular organisms rely on networks of metabolic interactions between cells and their environment. For microbial communities, the prediction of functions from community composition has recently become possible by learning statistical models from sparse experimental data sets, also enabling design of synthetic communities^1–3^. However, experimental studies have shown that abundances of metabolic pathways rather than of species are predictive of host-microbe metabolic interactions^4,5^. Experimental communities can also show identical functions with multiple stable compositions due to extensive metabolite cross-feeding^6^, and coexistence of species can be possible in multi-species communities but not in pairwise co-culture^7^. Pinpointing the largely unknown metabolic exchanges in such systems often relies on inference from *in vitro* experiments with simple consortia^8^. Thus, to understand and design metabolic interactions in complex communities in the context of ecosystem function, one should aim to characterize community metabolic functions comprehensively.

Metabolic pathway analysis aims for such comprehensiveness, using constraint-based models (CBMs) that are available for many organisms^9,10^. CBMs can represent the genome-scale metabolic network of an organism and allow for the analysis of metabolic fluxes (reaction rates). The common assumption of steady state results in a linear system of metabolite mass balances in which the variables are the fluxes, constrained by a stoichiometric matrix that represents the network structure, and any other linear constraints, such as upper and lower flux bounds. This defines a solution space containing all feasible combinations of steady-state fluxes, and each combination is known as a flux distribution^11^. The solution space is a flux cone if all constraints are homogeneous (i.e., if the mass balances always sum to zero), and a more general flux polyhedron if any of the constraints are inhomogeneous^12,13^.

In contrast to the commonly used flux balance analysis (FBA)^14^, which yields a single flux distribution that is not necessarily unique^15^, metabolic pathway analysis characterizes all network capabilities. Specifically, it defines the entire solution space in terms of fundamental and formally defined pathways such as elementary flux modes (EFMs)^16^. However, pathway counts grow combinatorially with network size^17,18^. This poses formidable challenges for the large-scale analysis of metabolic networks^19^. The main limitation is not necessarily the computation of pathways: state-of-the-art algorithms can enumerate all EFMs in networks with hundreds of reactions and billions of EFMs^20–23^. However, to obtain pathway sets amenable to analysis, one needs to limit enumeration to EFM subsets^24^. Examples include the shortest EFMs^25^, random EFMs^26^, EFMs involving specific reactions^27^, minimal sets of EFMs that generate the flux cone^28,29^, or EFMs that are consistent with thermodynamics^30^, regulation^31^, or metabolomics data^32^.

When addressing a specific biological question, complete enumeration may be achieved by using pathway definitions derived from EFMs that consider a sufficiently small subnetwork of interest within the full metabolic network. This amounts to projecting the flux space of the full network onto the sub-network, thus retaining all constraints on the full network^33^. Elementary conversion modes (ECMs) were originally defined as net metabolite conversions (i.e., for the subnetwork of reactions that exchange metabolites with the environment)^34,35^ but they could generalize to arbitrary subnetworks. One can further reduce pathway counts by ignoring the magnitude of fluxes and focusing on the unique flux patterns obtained by taking the signs of fluxes. For example, elementary flux patterns (EFPs) are flux patterns of EFMs projected onto a subnetwork, specifically those patterns that cannot be built from others without cancellation^36^.

Another key limitation of EFMs and pathway definitions derived from EFMs is that they are only defined for homogeneous constraints such as steady-state mass balances and reaction reversibilities. This means that they do not allow for important functional constraints such as experimentally determined bounds on growth or metabolite exchange rates^37^. Elementary flux vectors (EFVs) generalize EFMs by allowing arbitrary linear inhomogeneous constraints, but the challenges of enumeration remain^12,13,23^. Alternative approaches aim to find minimal reaction sets (MRSs)^38–40^ under inhomogeneous constraints, but these all rely on direct minimization of a mixed-integer linear program (MILP), which prevents scaling to large networks. Finding EFMs that satisfy inhomogeneous constraints poses the same challenges^41–43^.

To make pathway analysis more amenable to applications involving complex metabolic interactions in consortia and tissues, here we propose minimal pathways (MPs) as a generalization of existing pathway concepts that enables targeted analysis of subnetworks under inhomogeneous constraints. This makes MPs ideal for large-scale pathway analysis, and pathways where no reactions are dispensable are sufficient for key applications^13^. By developing efficient methods that find, sample, and enumerate MPs in large metabolic networks and subnetworks, we enable and demonstrate applications to bacterial central metabolism in a genome-scale context, host-microbe interactions in the human gut, and butyrate-producing microbial communities. These proofs of principle yield, for example, detailed predictions of biologically plausible metabolic exchanges between gut bacterial species and the human host. We envisage that MPs will be instrumental—along with statistical models and *in vitro* experimental data—in generating hypotheses on metabolic interactions in complex communities that can be tested *in vivo*.

## Results

### Minimal pathways are compact representations of metabolic capabilities

We define an MP as a minimal subset of reactions from a metabolic subnetwork, which in turn is a subset of reactions from a full metabolic network (see **Methods** for details). Specifically, we require *support-minimality*, meaning that all reactions in an MP need to be active (i.e., have non-zero flux) when no other reactions in the subnetwork are active in order to satisfy all constraints on the full network^13^. If no flux is required through the subnetwork, there are no MPs, so the constraints should include at least one functional requirement such as a minimal growth rate or flux. In other words, each MP represents a minimal way of performing one or more metabolic functions, and none of the reactions in that MP can be deactivated without disrupting those functions. The full set of MPs reveals reactions that never contribute to function (they do not participate in any MP), essential reactions (they participate in all MPs), and the relative functional importance (share of MPs in which they participate) of the remaining reactions.

Relative to existing pathway concepts, which differ in whether they generalize to subnetworks, allow inhomogeneous constraints, include flux stoichiometries or only patterns, and are support-minimal, MPs can be seen as a generalization (**Fig. 1A**). Specifically, combinations of fluxes in a subnetwork that satisfy all constraints on the full network lie in the projection of the full flux space onto the subnetwork, and the minimal set of vectors that generate this subspace without cancellations are its EFVs^12,13^. A subset of the EFVs must be support-minimal, and the flux patterns of these EFVs are equal to the set of MPs for a given subnetwork and network. Since EFVs are themselves a generalization of EFMs, this also implies that MPs are equal to the support-minimal flux patterns of EFMs and pathway definitions such as ECMs and EFPs, which are derived from EFMs, when all constraints are homogeneous. MPs also generalize MRSs from full metabolic networks to arbitrary subnetworks and from absolute to relative minimality (i.e., all MRSs have the same minimal number of reactions while some MPs can include more reactions).

**Figure 1.**
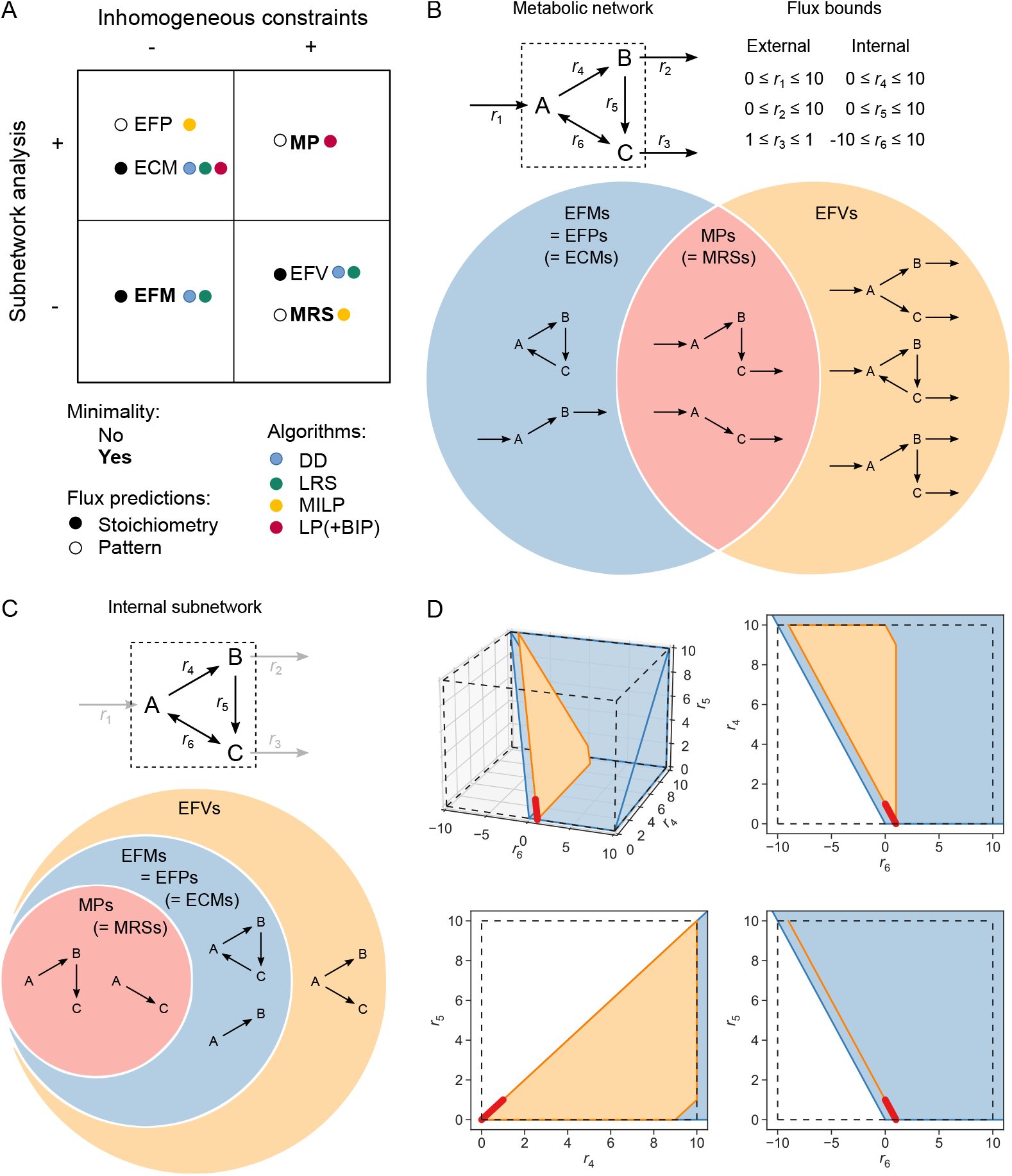
Comparison of metabolic pathway definitions. (**A**) Key features of elementary flux modes (EFMs), elementary flux vectors (EFVs), elementary flux patterns (EFPs), elementary conversion modes (ECMs), minimal reaction sets (MRSs), and minimal pathways (MPs). Pathway definitions are distinguished by whether they allow inhomogeneous constraints, enable targeted analysis of subnetworks, are support-minimal (bold text), and include flux stoichiometries (filled circles) or only patterns (open circles). Colored circles indicate the state-of-the-art algorithms for pathway enumeration: the double description method (DD), lexicographic reverse search (LRS), mixed-integer linear programming (MILP), and linear programming (LP) with or without binary integer programming (BIP). (**B**) Flux patterns of an example metabolic network. The network consists of three metabolites (A–C) participating in three boundary reactions that exchange metabolites with the environment (*r*_1_–*r*_3_) and three internal reactions (*r*_4_–*r*_6_). All reactions except *r*_3_ are irreversible, and we apply the given bounds to fluxes. The non-zero bounds are not taken into account by pathways corresponding to EFMs, which are not defined for inhomogeneous constraints. (**C**) Flux patterns for the subnetwork consisting of metabolites A–C and internal reactions *r*_4_–*r*_6_ within the full network in (B) with identical bounds. (**D**) Three-dimensional flux space and its two-dimensional projections corresponding to EFMs (blue), EFVs (orange), and MPs (red) for the subnetwork from (C) with bounds from (B).

To illustrate how pathway definitions relate to each other, we use an example metabolic network with six reactions and three metabolites (**Fig. 1B**). Counting pathways as flux patterns (i.e., considering flux signs but not stoichiometry) there are six EFVs, four EFMs, and three MPs. For flux patterns in full metabolic networks, EFMs are equal to EFPs. In principle, ECMs and MRSs could also be generalized to be equal to EFMs and MPs, respectively. The MPs correspond to the intersection of the EFMs and the EFVs, but note that EFMs do not take inhomogeneous flux bounds into account. Also, this network allows for a thermodynamically infeasible loop, which is avoided by the MPs.

Next, consider the subnetwork consisting of the three internal reactions and metabolites from the full metabolic network (**Fig. 1C**), there are two flux patterns for MPs, four for EFMs, and five for EFVs. Again, the EFMs are equal to the EFPs, and the ECMs and MRSs could generalize to EFMs and MPs, respectively. The MPs are a subset of the EFMs, which in turn are a subset of the EFVs. We also saw the same patterns for the subnetwork consisting of the three external reactions (**Fig. S1**). Thus, for both full networks and subnetworks, MPs are compact representations of metabolic capabilities that form a subset of other pathway definitions, and this is also apparent from the corresponding feasible flux spaces (**Fig. 1D**). This illustrates the generalization by MPs, namely to yield minimal functional representations of subnetwork fluxes while accounting for all of the full network’s constraints.

### Iterative minimization and pathway graphs find minimal pathways efficiently

The different formal definitions of metabolic pathways also entail different algorithmic options—and limitations—for pathway enumeration (see **Fig. 1A** for an overview). EFMs are usually enumerated by derivatives of the double description (DD) method, which scales poorly with network size^20,22^. However, it has recently been shown that lexicographic reverse search (LRS) can perform as well as DD or better^23^. EFVs and ECMs are also enumerated by DD^13,35^ or LRS^23,44^. EFPs and MRSs both rely on MILPs for enumeration, which is problematic for scaling^36,40^. ECMs can additionally be enumerated by solving many parallelizable linear programs (LPs), and individual LPs can be solved efficiently^35^.

To find MPs efficiently, we developed an iterative minimization approach that leverages support-minimality: to find an MP, we iteratively remove reactions from the subnetwork and minimize subnetwork flux with an LP until no further reactions can be removed without violating any of the constraints on the network as a whole (**Algorithm S1**). A randomized version allows random sampling of MPs. To enumerate MPs, we alternate iterative minimization with computation of minimal cut sets (minimal sets of reactions that block network functions when deactivated; MCSs) in a separate binary integer program (BIP) (**Algorithm S2**).

This is similar to an approach suggested for EFMs^45^, which approximates EFMs with a single LP and therefore does not guarantee support-minimality. We also realized that maximal cliques (i.e., fully connected subgraphs) in a simple graph representation of known MPs can predict subnetworks likely to contain unknown MPs. This drastically reduced the sizes of optimization problems for finding MPs and lead us to the graph-based iterative minimization algorithm for MP enumeration (**Algorithm S3**).

To illustrate the different approaches for MP enumeration and sampling, we use a random subnetwork of 562 reactions in the genome-scale *Escherichia coli* model iML1515^46^ with 2,868 irreversible reactions. Graph-based enumeration found all 27,648 MPs within 45 minutes (**Fig. 2A**), but after running for one hour, the other methods returned an order of magnitude fewer MPs (**Fig. 2B**). By design, iterative minimization also found all MCSs in parallel with the MPs (**Fig. 2C**). In more detail, direct minimization only identified MPs of the minimal size 14, iterative minimization without graph and randomization found MPs in arbitrary, but clearly not random order, and randomized iterative minimization found MPs in apparently random order (**Fig. 2D**). The order of MP sizes for graph-based enumeration was more structured than for the other iterative approaches, with a tendency to first find small MPs and to find many MPs of similar size after one another. The effective subnetwork size in graph-based enumeration varied between the full subnetwork when no new cliques were available, and much smaller sizes approximately equal to the sizes of MPs when cliques were available (**Fig. 2E**). These results clearly favor all variants of iterative minimization over direct minimization.

**Figure 2.**
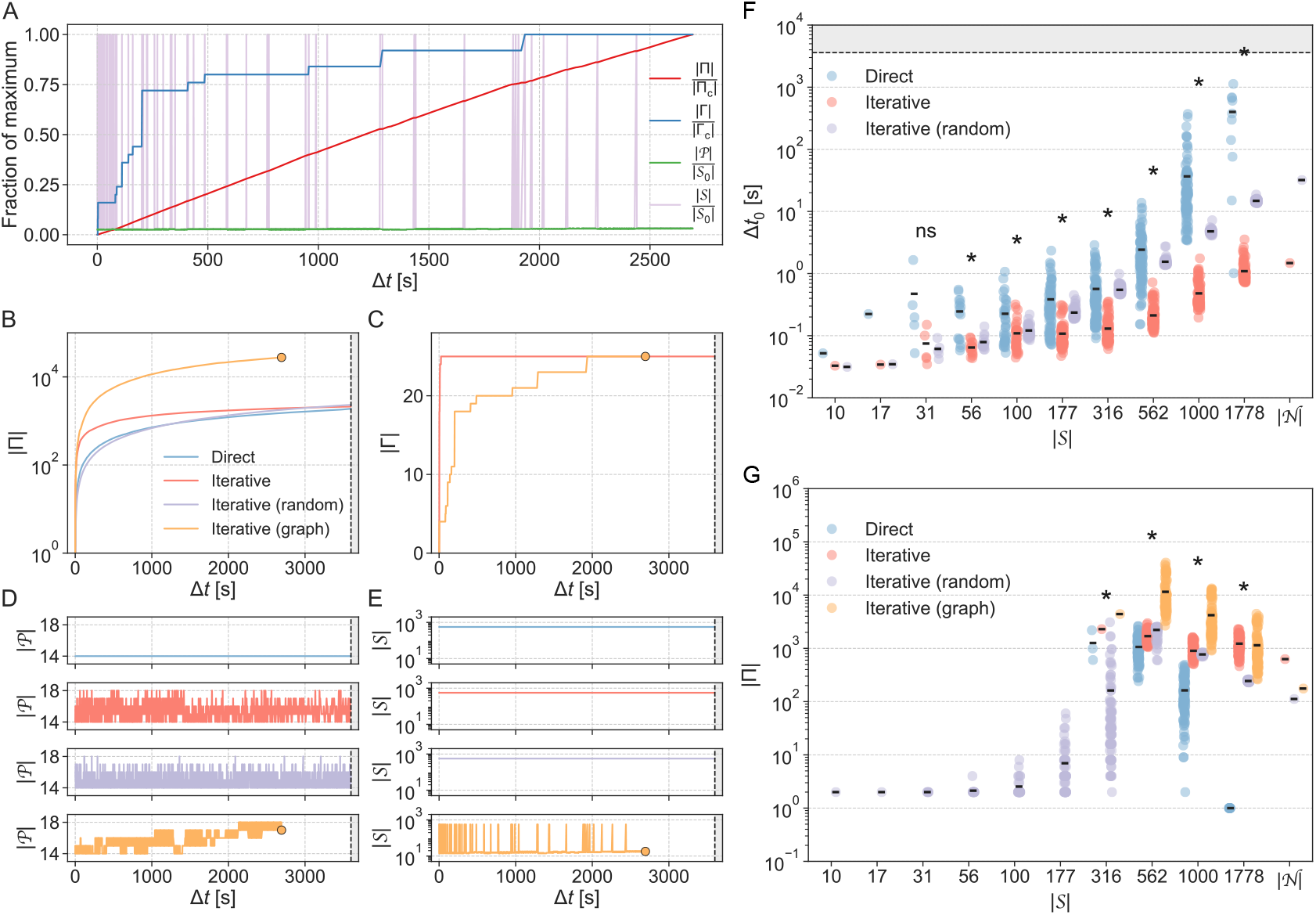
Benchmarking of MP enumeration and sampling methods for *Escherichia coli*. (**A**) Complete enumeration of MPs and MCSs using iterative minimization with graph in a random subnetwork of size | *𝒮*_0_ | = 562 within the genome-scale *E. coli* model iML1515 with | *𝒩* |= 2,868 irreversible reactions. The number of MPs | Π | and MCSs | Γ| found is shown along with the MP size | *𝒫* | and effective subnetwork size | *𝒮* | as a function of running time Δ*t*. Values are normalized by the size of the complete set of MPs | Π_c_ |, the size of the complete set of MCSs | Γ_c_|, or the initial subnetwork size | *𝒮*_0_ |. (**B-E**) Comparison to incomplete enumerations in the same subnetwork performed with direct minimization and iterative minimization with and without randomization, shown separately for |Π | (**B**), | Γ | (**C**), | *𝒫* | (**D**), and | *𝒮* | (**E**). End points of the completed enumeration are indicated by dots. (**F**) Running time Δ*t*_0_ for finding the first new MP with direct (blue) and iterative minimization with (purple) and without (red) randomization across subnetwork sizes | 𝒮 | in iML1515. Stars indicate significant difference between means from one-way ANOVA where data was available for all three methods, or *t*-test where data was available for two methods (significance level 0.05). Computation was stopped after one hour (indicated by dashed line). (**G**) Number of MPs | Π | found in one hour with direct minimization (blue), iterative minimization (red), iterative minimization with randomization (purple), and iterative minimization with graph (orange) across subnetwork sizes | *𝒮* | in iML1515. Enumerations that were completed within one hour are not included. Statistics as in (F).

For a more systematic assessment, we varied subnetwork sizes by sampling 100 subnetworks each of sizes ranging from 10 reactions to the full network (**Supplementary Data 1**) in six models of microbial and mammalian cells ranging in size from about 100 to more than 14,000 irreversible reactions (**Table S1**). The running time to find the first MP was almost always greater and grew faster with subnetwork size for direct than for our different variants of iterative minimization (**Fig. 3A–C** and **Fig. S2**). Because running times also depend on the size of the full network, we used this size for normalization to compare between models. Normalized running times for finding the first MP scaled similarly between models, and the scaling was well described by the double exponential model log *y* = *a*+*b*^*x*^ (**Fig. 3D**). We estimated the scaling parameters as *b* = 2.3±0.02 for iterative minimization, *b* = 4.0±0.06 for randomized iterative minimization, and *b* = 7.3± 0.13 for direct minimization, revealing large differences in scaling between methods. This is mirrored by the number of MPs found in incomplete enumerations when we restricted the running time to one hour, demonstrating superior performance of graph-based enumeration—which found an order of magnitude more MPs than the other methods in most subnetworks—in particular (**Fig. 3E–G** and **Fig. S3**). Altogether, our algorithms using iterative minimization, especially when accelerated by pathway graphs, systematically outperformed the state-of-the-art direct minimization algorithm in terms of speed and scalability.

**Figure 3.**
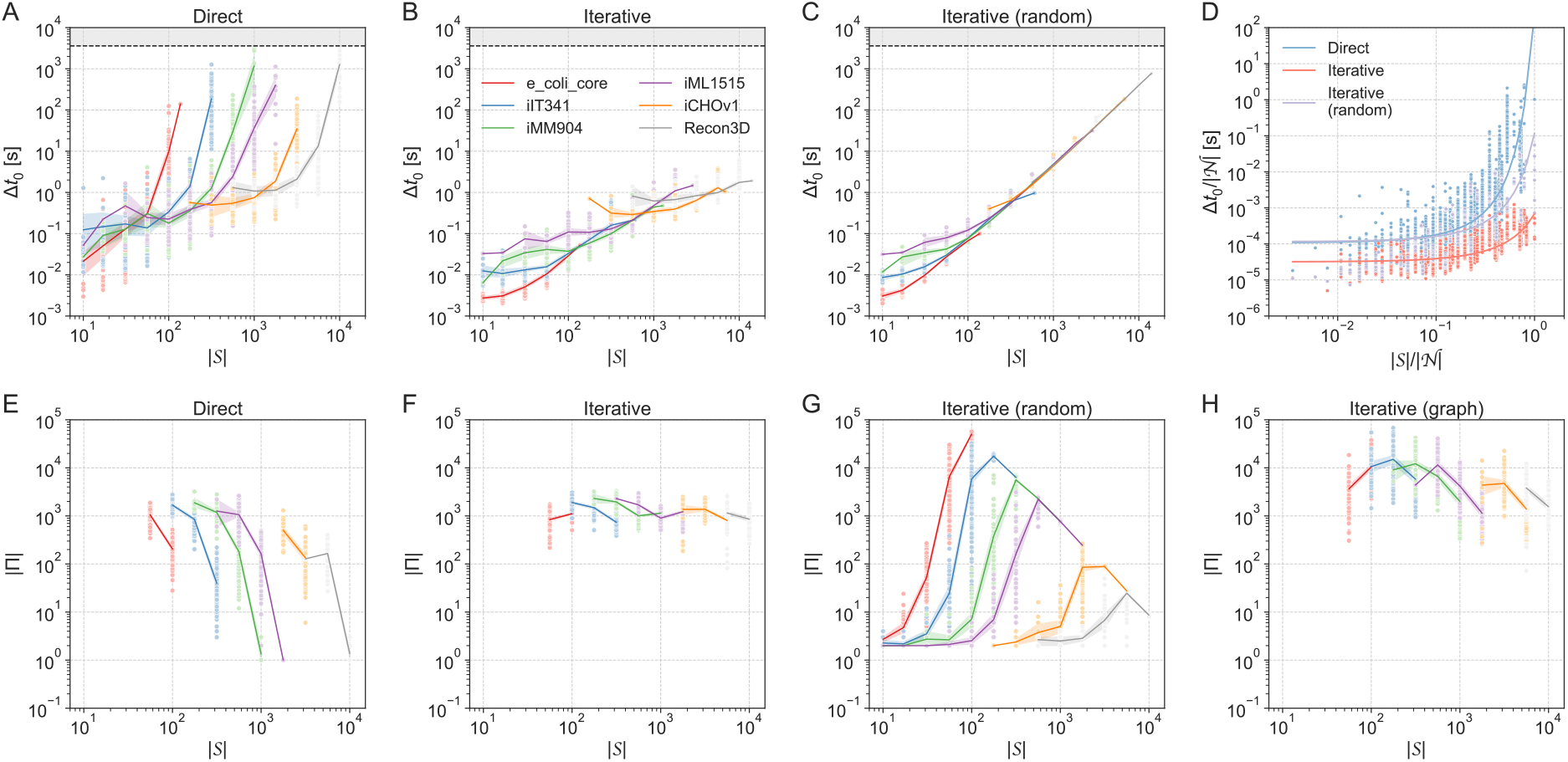
Benchmarking of MP enumeration and sampling methods for a range of microbes and mammals. (**A–C**) Running time Δ*t* _0_ as a function of subnetwork size | *𝒮* | for (**A**) direct minimization, (**B**) iterative minimization, and (**C**) randomized iterative minimization. Enumeration was stopped after one hour (indicated by dashed line). (**D**) Running time Δ*t*_0_ as a function of subnetwork size | *𝒮* |, both normalized by network size | *𝒩* |, for direct (blue), iterative (red), and randomized iterative (purple) minimization. Lines are fits of the double exponential model log_10_ (Δ*t*_0_/|*𝒩* | = *a* + *b*^| *𝒮* | /| *𝒩* |^. (**E-H**) Number of MPs | Π | found in one hour as a function of subnetwork size *𝒮* for (**E**) direct minimization, (**F**) iterative minimization, (**G**) randomized iterative minimization, and (**H**) iterative minimization with graph. Enumerations that were completed within one hour are not included. Lines indicate means with 95% confidence intervals from bootstrapping and colors indicate models.

### Minimal pathways enable comprehensive analysis of large-scale metabolic systems

Scalability of pathway analysis requires that pathway sets are small enough to be analyzed meaningfully with respect to the biological problem addressed, while still characterizing the full flux space relevant for this problem. MPs minimize the number of pathways to enumerate and analyze, but for all six example models we used for benchmarking, the number of MPs found in complete enumerations still increased exponentially with subnetwork size (**Fig. 4A**). However, between the models, which vary by two orders of magnitude in terms of number of reactions, we found neither an exponential increase of MP set size with subnetwork size nor a clear correlation between the two.

**Figure 4.**
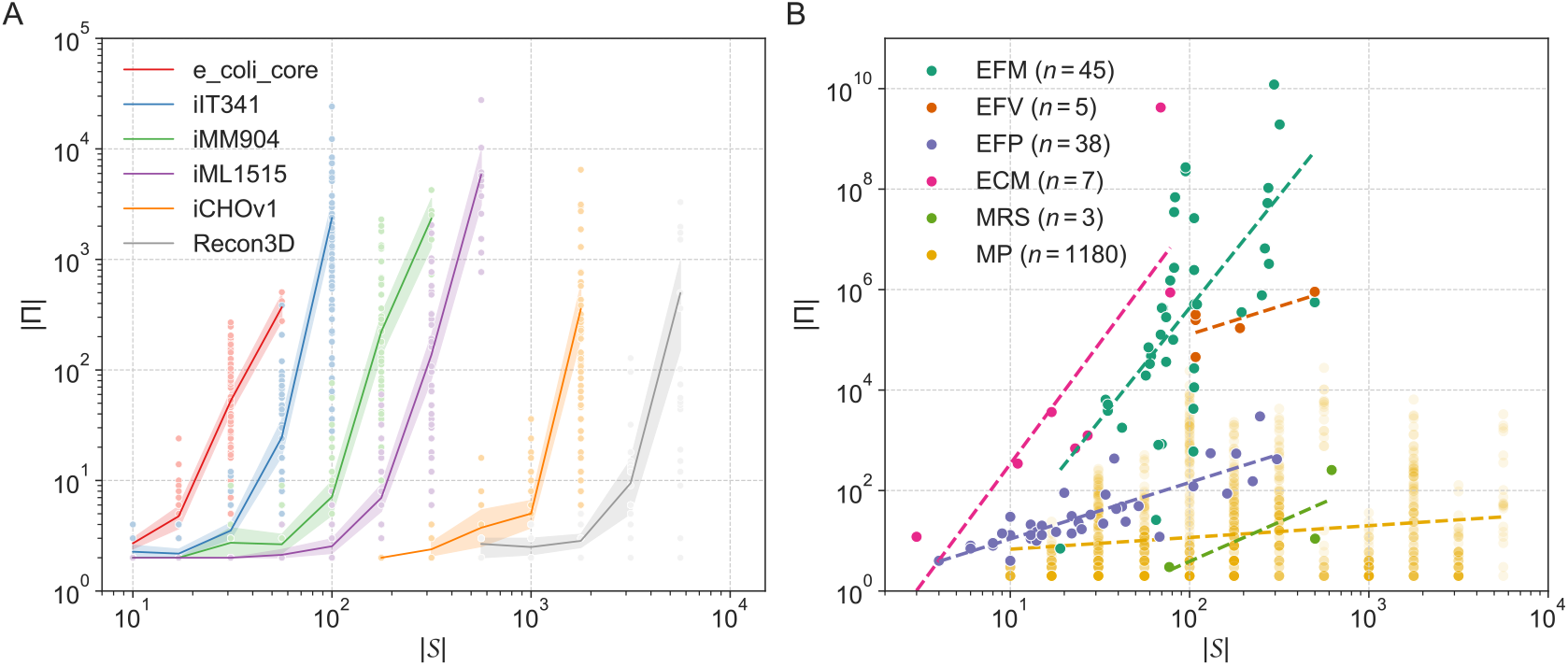
Pathway counts as a function of metabolic subnetwork size. (**A**) Number of MPs | Π | as a function of subnetwork size | *𝒮* | for complete enumerations of MPs in six microbial and mammalian models. Lines indicate means with 95% confidence intervals from bootstrapping and colors indicate models. (**B**) Number of MPs | Π | as a function of subnetwork size | *𝒮* | for all the complete enumerations of MPs shown in (A) and complete enumerations of EFMs, EFVs, EFPs, ECMs, and MRSs in 20 previous studies. For EFMs, EFVs, and MRSs, subnetwork size is the same as the full network size. Dashed lines show linear fits for the individual pathway definitions.

To relate MPs to other pathway concepts, we compared MP counts for all benchmarking enumerations that were completed within one hour to results for 98 networks from 22 published studies reporting complete enumeration of EFMs, EFVs, EFPs, ECMs, or MRSs (**Fig. 4B** and **Supplementary Data 2**). MPs have been enumerated in larger networks and subnetworks than other pathways, up to an order of magnitude. More importantly, MP numbers scale better with network and subnetwork size than any other pathway concept, with the possible exception of MRSs, which are closely related to MPs but have few available data points. This empirical evidence supports the theoretical argument that MPs provide compact representations of metabolic network capabilities relative to other pathway concepts, which should make MPs most suitable for large-scale applications.

### Sampled pathways predict growth effects of gene knockouts in *E. coli* central metabolism

To evaluate the application potential of MPs, we first predicted growth effects of gene knockouts, which is a standard test scenario for CBMs. Gene essentiality is also increasingly recognized as quantitative and context-dependent^47^. Even for model organisms such as *E. coli*, it is still unclear what the essential genome is, with targeted knockout libraries^48,49^, randomized transposon mutagenesis^50^, and most recently CRISPRi screens^51,52^ yielding consensus as well as divergent gene sets. To capture gene essentiality quantitatively, we applied our randomized iterative minimization procedure to the central metabolism of *E. coli* as a subnetwork in the genome-scale model iJO1366^53^. This subnetwork of 153 irreversible reactions, mostly from glycolysis, the tricarboxylic acid (TCA) cycle, and the pentose phosphate pathway (**Fig. 5A**), is embedded in the full model of 3,219 irreversible reactions.

**Figure 5.**
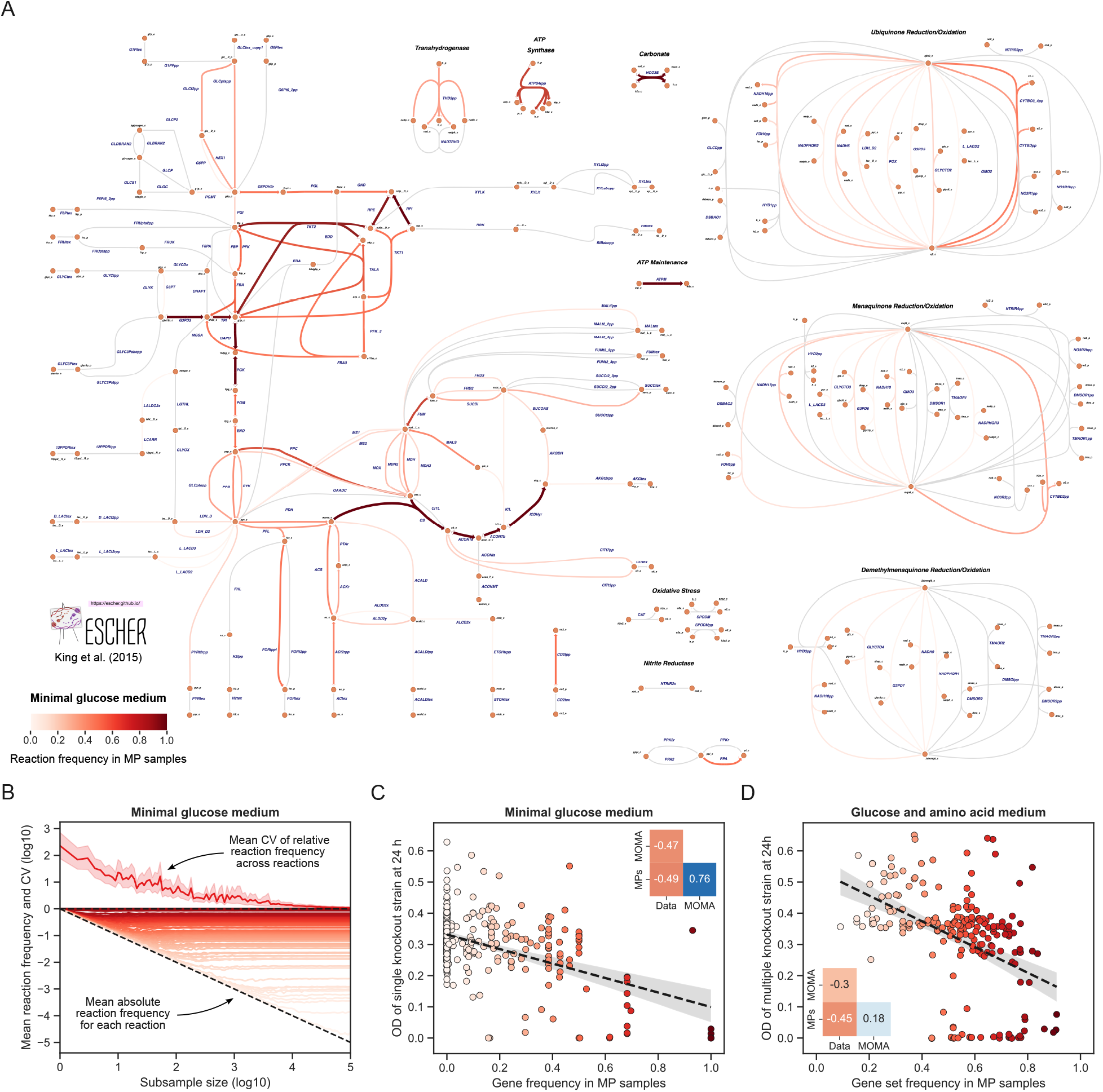
MP sampling of *E. coli* central carbon metabolism. (**A**) Pathway map of *E. coli* central carbon metabolism made with Escher 54. Arrow color and width indicate reaction frequency across 100,000 randomly sampled MPs. The network was used as a subnetwork within the genome-scale model iJO1366 with aerobic conditions, a minimal glucose medium, and growth and ATP maintenance requirements. For reversible reactions, frequency does not distinguish between flux directions. (**B**) Convergence of reaction frequencies in MP subsamples. The 100,000 MP samples were subsampled with replacement 100 times for 100 logarithmically spaced subsample sizes. Mean absolute reaction frequency for each reaction is shown below, and mean coefficient of variation (CV) of relative reaction frequency across all reactions is shown above with 95% confidence interval. Dashed lines indicate *y* = 0 and *y* = −*x*, the line along which reaction frequencies converge with increasing subsample size. Colors indicate frequency as in (A). (**C**) Optical density (OD) of single gene knockout strains at 24 h as a function of gene frequency in MP samples for a minimal glucose medium. Colors indicate frequency as in (A). The dashed line shows a linear fit with 95% confidence band, and the inset shows Pearson correlations between data, MP gene frequencies, and growth predicted by minimization of metabolic adjustment (MOMA). (**D**) OD of multiple gene knockout strains at 24 h as a function of gene set frequency in MP samples for a glucose and amino acid medium, otherwise as in (C).

To predict the relative importance of enzymes in central carbon metabolism for aerobic growth on a minimal glucose medium, we randomly sampled 100,000 MPs and calculated the reaction frequencies (i.e., the fraction of MPs in which each reaction participated; **Supplementary Data 3**). We confirmed that the sample size was large enough for convergence of reaction frequencies; on average, the frequencies converged linearly with increasing subsample size (**Fig. 5B**). No reactions in the pentose phosphate pathway were essential, but five enzymes participated in more than 80% of all MPs (transaldolase, transketolases, ribulose 5-phosphate 3-epimerase, and ribose-5-phosphate isomerase). Also, four enzymes stood out in the TCA cycle (citrate synthase, aconitase, isocitrate dehydrogenase, and fumarase), appearing to be much more important for aerobic growth on glucose than, for example, 2-oxogluterate dehydrogenase and succinyl-CoA synthetase. As expected, isozymes such as the malate dehydrogenases in the TCA cycle had similar and intermediate frequencies. These patterns are consistent with the alternative operation of the full TCA cycle and the PEP-glyoxylate cycle for complete glucose oxidization^55^.

To evaluate our predictions quantitatively, we mapped samples to enzyme-encoding genes and compared gene frequencies across MPs to experimental growth data for single and multiple gene knockout strains (**Supplementary Data 3**). Gene frequencies in sampled MPs and growth effect predictions from minimization of metabolic adjustment (MOMA)^56^ correlated equally well with growth of single knockout strains^48^ as well as strongly with each other (**Fig. 5C**). For multiple knockout strains on a non-minimal glucose and amino acid medium^49^, MP predictions maintained their quality, but MOMA predictions deteriorated, and the strong correlation between MPs and MOMA disappeared (**Fig. 5D** and **Fig. S5**). More specifically, a few genes with large residuals primarily drove deviations between MP predictions and growth data (**Fig. S6** and **Fig. S7**). MOMA predictions generally tended to underestimate knockout effects on growth, in particular for the non-minimal glucose and amino acid medium (**Fig. S8**). Overall, predictably converging sampling of MPs to compute reaction and gene frequencies in an unbiased manner may be more suitable for predicting context-dependent growth effects of gene knockouts than MOMA, the biased reference method.

### Metabolite exchanges explain balanced growth in a host-microbe model of the human gut

To demonstrate that MP sampling and enumeration can help address biological questions in large communities with multiple interacting species, we applied it to a host-microbe model of the human gut. The study establishing this model predicted potential metabolic exchanges with FBA^57^. Here, we aimed to identify key metabolic cross-feedings and used updated models of all species^58,59^ (**Table S2**). In our host-microbe model, a human cell interacts with six commensal microbes in a rich metabolic environment with aerobic conditions for the human cell and anaerobic conditions for the microbes. The organisms can exchange metabolites with the environment through external exchange reactions and with each other through internal exchange reactions to a shared compartment (**Fig. 6A**). To simulate cross-feeding between the microbes and from the microbes to the host, the microbes are allowed to consume and produce shared metabolites, while the human cell can only consume shared metabolites.

**Figure 6.**
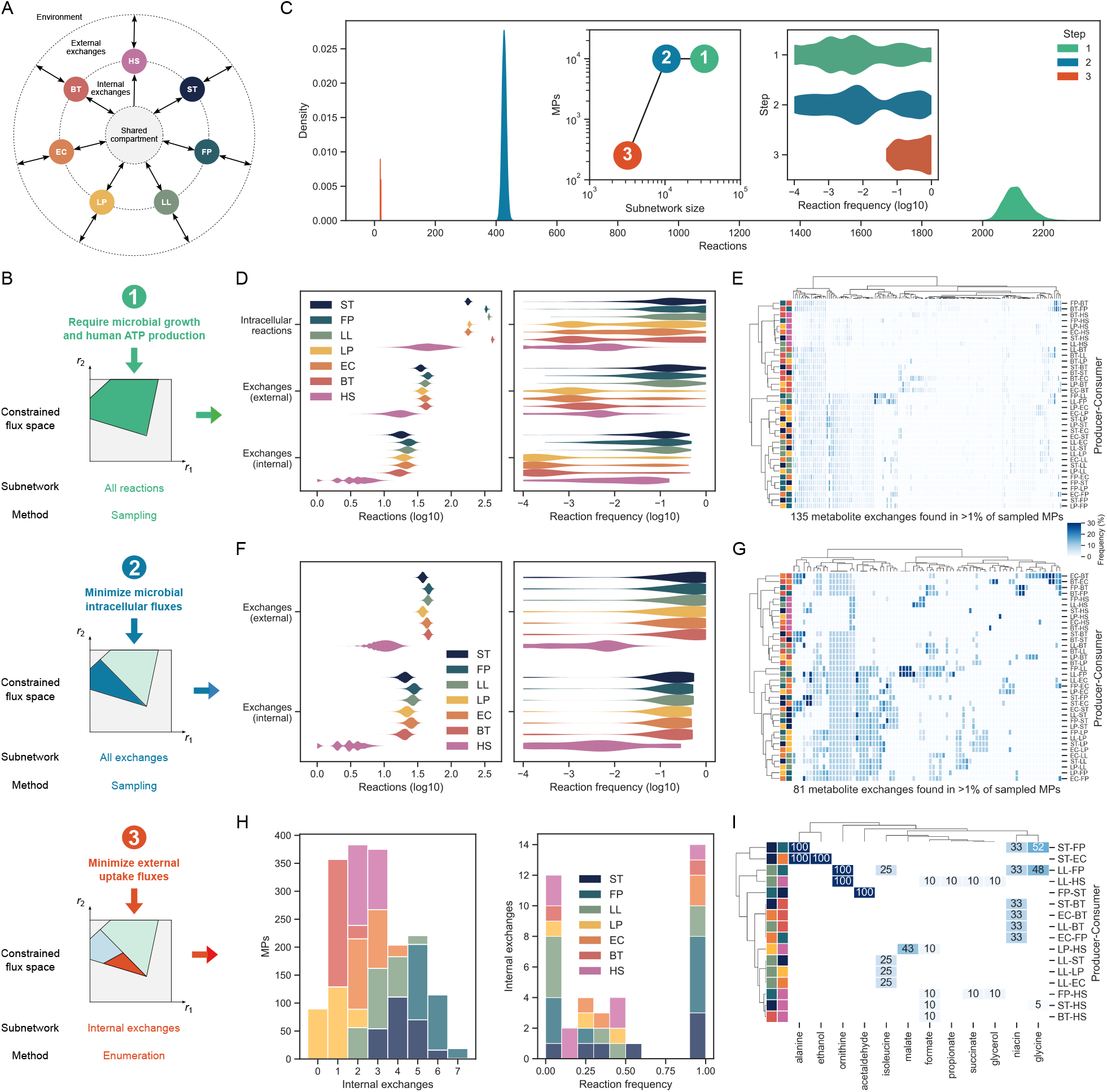
MP sampling and enumeration of a sequentially constrained host-microbe model of the human gut. (**A**) Host-microbe model structure with organisms connected to each other through internal exchanges and to the environment through external exchanges. Organisms: *Streptococcus thermophilus* (ST), *Faecalibacterium prausnitzii* (FP), *Lactococcus lactis* (LL), *Lactobacillus plantarum* (LP), *Escherichia coli* (EC), *Bacteroides thetaiotaomicron* (BT), and *Homo sapiens* (HS). (**B**) The host-microbe model was constrained sequentially by requiring (1) balanced growth for the six microbes and ATP production for the host, (2) minimal intracellular flux for the microbes, and (3) minimal external exchange flux with the environment. We found MPs after each step by (1) sampling the full network, (2) sampling the subnetwork of exchanges, and (3) enumerating the subnetwork of internal exchanges. (**C**) Reactions in sampled and enumerated MPs for each step. The insets show sampled or enumerated MPs by subnetwork size and reaction frequency for each step. (**D–I**) Reactions in MPs and their frequency across MPs by reaction type and organism (left) and frequency of metabolite exchanges between organisms (right) after the first (**D–E**), second (**F–G**), and third (**H–I**) step. Distributions are kernel density estimates cut at the extreme observations. In the heatmaps, rows are producer-consumer pairs, columns are exchanged metabolites, and each cell indicates metabolite exchange frequency across MPs. Rows and columns are clustered by Ward’s minimum variance method.

To identify increasingly specific and important sets of cross-fed metabolites that could explain human energy generation and balanced microbial growth in the gut, we applied constraints sequentially to the model^37^ (**Fig. 6B**). Aiming to explain as much of these functional requirements as possible in terms of metabolite cross-feeding with biologically reasonable microbial flux distributions^60,61^, we required (1) balanced growth for the microbes and ATP production for the human cell, (2) minimal microbial intra-cellular flux, and (3) minimal metabolite exchange flux with the environment. We first sampled 100,000 MPs in the full network (33,272 reactions), then sampled 100,000 MPs in the subnetwork of external and internal exchanges (10,452 reactions), and finally enumerated all 252 MPs for the subnetwork of internal exchanges (3,185 reactions). Reaction frequencies converged as expected for the samples and graph-based enumeration only took about 33 minutes on a laptop (**Fig. S9**).

Adding constraints and reducing the size of the subnetwork with each step reduced the number of reactions in each MP, while the frequencies of reactions across MPs increased (**Fig. 6C** and **Supplementary Data 4**). In particular, internal metabolite exchanges became fewer and more frequent after each step, providing increasingly specific predictions. The first step found 3,031 exchanges of 447 metabolites, but none that were very frequent, especially not for large microbes such as *Bacteroides thetaiotaomicron* with more metabolic capabilities (**Fig. 6D–E**). The second step reduced this to 1,027 exchanges of 98 metabolites with higher frequencies that were also much more similar between microbes thanks to the requirement of minimal intracellular flux (**Fig. 6F–G**). The set was finally narrowed down to 30 exchanges, six of which were essential, of 12 metabolites, including amino acids, fermentation products, TCA intermediates, and the B vitamin niacin (**Fig. 6H–I**). Specifically, we predicted intermicrobial cross-feeding of acetaldehyde, alanine, ethanol, glycine, isoleucine, niacin, and ornithine, while the human cell was predicted to consume microbial formate, glycerol, glycine, malate, ornithine, propionate, and succinate.

Importantly, the predicted cross-fed metabolites have been observed in the gut and implicated in the maintenance of host-microbiota homeostasis as well as human health and disease. For example, a possible alanine exchange predicted by the same microbial model collection was confirmed *in vitro*^58^, increased isoleucine levels have been associated with improved growth in humans^62^. Consumption of glycine by the microbiota has been shown to reduce glycine levels in the host, which has been associated with non-alcoholic fatty liver disease (NAFLD), obesity, and type 2 diabetes^63^. The gut microbiota also contributes more directly to NAFLD by producing ethanol^64^ and byproducts of its degradation such as acetaldehyde^65^. Glycerol has a large influence on metabolism and composition of human gut microbial communities^66^, elevated levels and cross-feeding of microbially derived formate have been associated with inflammation^67^, and levels of succinate as well as microbes producing or consuming it have been associated with obesity and related diseases^68^. Exchange of niacin (B3) and its importance for microbiota stability are supported by *in silico, in vitro*, and *in vivo* evidence^4,69,70^, and microbial ornithine production promotes healthy gut mucosa through crosstalk with the host^71^. Finally, propionate is a short-chain fatty acid that, along with butyrate, is known to be cross-fed from gut bacteria to the host with beneficial effects on human health^72^.

In summary, MP sampling and enumeration allowed detailed analysis of a very large multicellular model, including prediction of metabolite exchanges between organisms that could explain growth and energy generation in the gut. Cross-feeding predictions appear plausible and provide hypotheses for specific producers and consumers, which are hard to identify experimentally.

### Pathway enumeration enables design of butyrate-producing microbial communities

A further application in multicellular systems, which requires both scalable algorithms and a flexible pathway definition, is design of communities by sampling or enumerating minimal sets of organisms capable of performing specific tasks in a given environment. To evaluate the potential of MPs for this purpose, we focused on synthetic microbial communities that produce butyrate, a short-chain fatty acid relevant for gut health^72^. Using design-experiment-model cycles with statistical models, Clark et al.^73^ explored and predicted how well communities consisting of subsets of 25 common gut bacterial strains could produce butyrate, revealing non-monotonic dependencies of butyrate production both on the overall number of species, and on the prevalence of the five known butyrate-producing strains. Using the medium and strain catalog from Clark et al.^73^ and genome-scale models of the 25 microbes^58^ (**Table S3**), we combined the models into a community model with metabolite exchange through a shared compartment. We then tested how well the model predicted butyrate production in 790 experimentally tested communities^73^. Requiring each strain to grow efficiently at 80% of its maximal growth rate per total sum of intracellular fluxes (a single insensitive parameter estimated from the data), predicted and experimental values correlated well (Pearson’s *r* = 0.73, weighted by number of measurements; **Fig. S10**), a prediction quality comparable to an earlier model fitted on data for 156 communities^73^.

Next, we used MP enumeration to predict species-minimal communities and their cross-feeding interactions while requiring efficient growth (80%) and at least 50%, 75%, 90%, or 95% of the maximal butyrate production of the complete 25-strain community (**Fig. 7A, Fig. S11**, and **Supplementary Data 5**). Note that increasing the maximal butyrate production of a community can be achieved trivially by adding more strains; MPs can find high-performance communities with minimal complexity (i.e., number of participating strains). As expected, the number of minimal communities tended to decrease with increasing production requirements (**Fig. 7B**). Finding more communities for 95% than for 90% is a consequence of minimality: all six communities for 95% were feasible, but not minimal for 90%. Typically, communities with five to ten members were needed for high butyrate production. This means that, apart from at least one of the five butyrate producers, minimal communities needed supporting strains to cross-feed intermediate metabolites to the butyrate producers (**Fig. 7C**). We also tested whether equally well-performing communities could be composed randomly by combining butyrate producers with random non-producers, keeping the same size and butyrate producer distributions as in the minimal communities (**Methods**). The substantial decrease of random community performance highlights the impact of specific community interactions on butyrate production capabilities (**Fig. 7D**).

**Figure 7.**
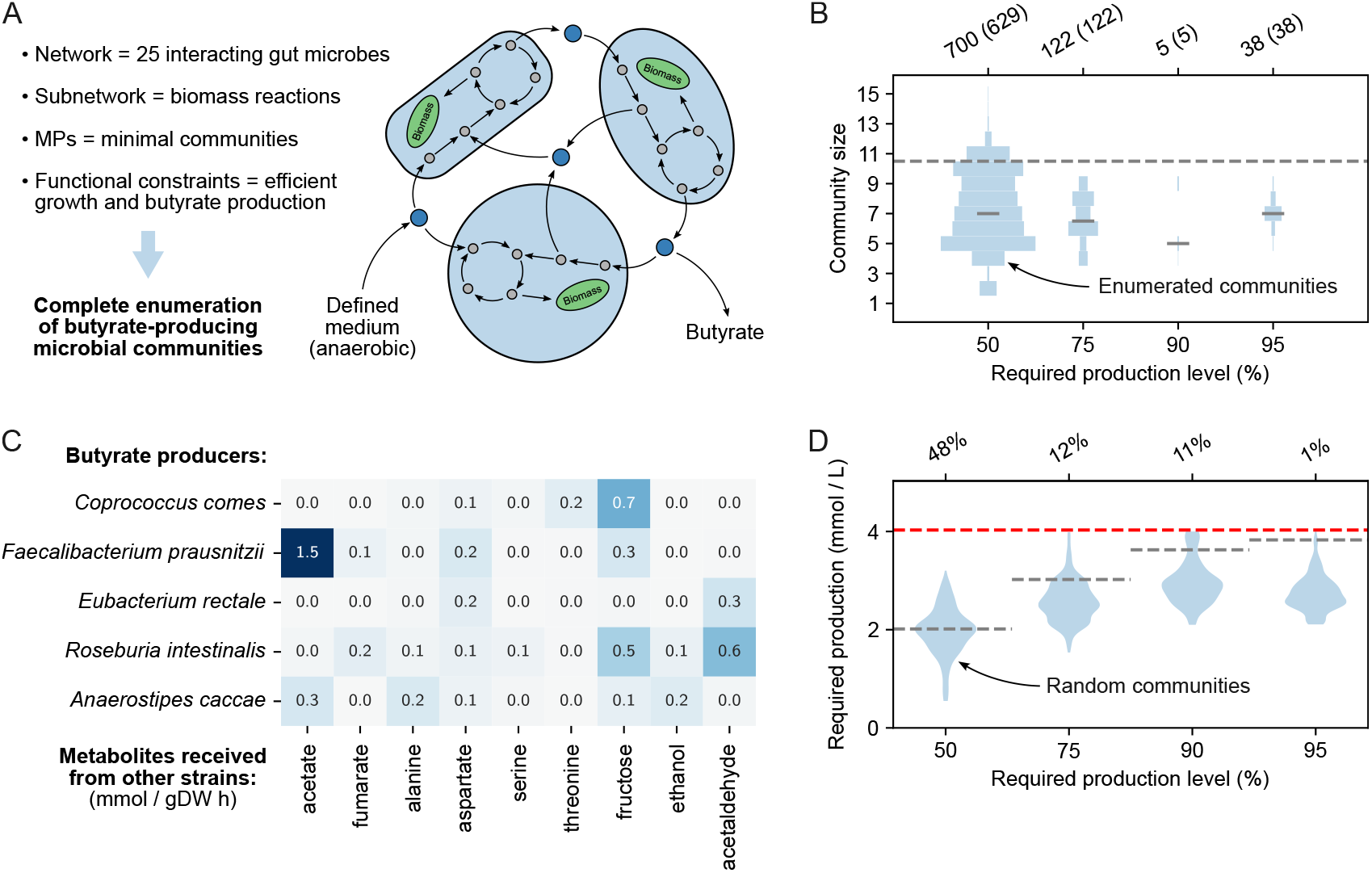
Enumeration of minimal gut communities for butyrate production. (**A**) Illustration of setup and assumptions for the enumeration of minimal butyrate-producing communities. (**B**) Size distributions of enumerated communities for increasing butyrate production requirements relative to the maximum of the full community. The total number of MPs (and of MPs with ten members or less) is indicated above. The area of each histogram is proportional to the number of MPs. (**C**) The quantitatively most important metabolites received by the butyrate producers via cross-feeding from other species for the communities enumerated at 50% required butyrate production. Rows are organisms, columns are metabolites, and each cell indicates the average cross-fed amount of each metabolite (mmol gDW^−1^h^−1^) received by a butyrate producer. (**D**) Butyrate production of random communities (500 per violin plot) constructed with the same butyrate producers and size distributions as the enumerated communities in (**A**). The red dashed line indicates the maximum butyrate production of the full community and the grey dashed lines indicate the required butyrate production level of the enumerated communities. The percentage of random communities achieving the required butyrate production level is indicated above.

To elucidate the relevant community interactions, we identified the top metabolites cross-fed to each butyrate producer in terms of average cross-fed quantity from other strains (**Fig. 7C** and **Supplementary Data 6**). Most of the exchanged metabolites were predicted to be important for multiple strains, consistent with expected benefits for both producer and consumer. Also, similar to our host-microbe predictions, predicted metabolite exchanges were supported by *in silico, in vitro* or *in vivo* evidence. Acetate, fumarate, and lactate are commonly secreted anaerobic fermentation products of gut microbes that are relevant for ATP production and redox balance. Acetate and fumarate consumption by the butyrate producer *Faecalibacterium prausnitzi* is corroborated experimentally^74,75^. The amino acids alanine, aspartate, glutamate, and threonine are beneficial for the consumers as building blocks of peptides, and microorganisms commonly leak amino acids^76^. As a sugar, fructose is beneficial for consumption in most microbes. However, *Bacteroides uniformis* has a pathway that uses arabinose from the medium to produce and secrete fructose, converting one NADH to NAD in the process. Ethanol and acetaldehyde are microbial fermentation products with inflammatory impact on the gut^77^. Though not experimentally validated, genomic evidence of ethanol and acetaldehyde consumption capabilities in gut microbes such as *Roseburia intestinalis* is interesting, as administration of *R. intestinalis* is known to ameliorate alcoholic fatty liver^78^. Hence, large-scale application of MPs to design of minimal butyrate-producing communities of gut microbes yielded hypotheses both on far-from-random high-performance communities and on their underlying metabolic interactions.

## Discussion

Pathway analysis for characterizing potentially complex metabolic interactions comprehensively in applications usually starts with a targeted biological question and prior knowledge or experimental data. Our generalization of pathway concepts to MPs addresses these two aspects by allowing for arbitrary subnetworks and by accounting for all linear constraints on the network as a whole, including inhomogeneous constraints. The definition of MPs as the support-minimal flux patterns of EFVs makes it straight-forward to reconcile MPs with available theory: MPs correspond to a subset of the EFVs, leading to compact pathway sets. Minimality implies that all reactions in an MP contribute to fulfilling functional requirements as defined by constraints on the full network, thereby connecting predictions directly to function and avoiding the pervasive problem of thermodynamically infeasible loops^79^. By design, MPs do not account for all possible flux distributions of the subnetwork, but rather the minimal flux distributions that support function. This is unlikely to limit MP applications because such minimal flux distributions are consistent with omics data^60,61^. To address the limitation of MPs being flux patterns without stoichiometry, one can apply FBA and related methods to computed MPs.

Efficient and scalable computations are a prerequisite for any pathway analysis of sufficiently complex organisms or communities. In general, we argue that our iterative minimization approach based on LPs can replace the computationally hard direct minimization of an MILP, which is the state of the art for pathway enumeration and sampling^26,36,41^. Through substantially improved scaling of running time, iterative minimization allowed MPs to be enumerated or sampled in networks of all sizes. Future algorithmic developments could focus on two aspects. First, in graph-based minimization, we detected all maximal cliques at every graph update, that is, when the full subnetwork was used to find an MP. This could be made more efficient by directly finding cliques with edges between reactions in the latest MP. Second, the main bottleneck for MP enumeration is the BIP used to find cut sets, which slows down each iteration, especially as the number of known MPs and MCSs increases. A faster procedure for finding new cut sets could substantially improve scaling, for example, by leveraging the duality of pathways and cut sets^80^.

To highlight different facets of the concept and computation methods for MPs, we selected three applications as examples. First, random sampling of MPs in *E. coli* central carbon metabolism within a genome-scale network demonstrated the use of a subnetwork to address specific biological questions such as quantitative gene essentiality. Importantly, MP-predicted relative importance of reactions and enzyme-encoding genes for growth agreed better with experimental gene knockout data than the standard MOMA method. Second, we sequentially constrained and identified MPs in a host-microbe model of a human cell interacting with six gut microbes. This shows that MP sampling can be used to analyze the full metabolic networks of multicellular systems and that adding constraints and reducing subnetwork size can make MPs enumerable even in some of the largest models considered to date. Cross-feeding predictions included metabolites with experimental evidence and possible roles in gut homeostasis and human health. Finally, we enumerated minimal microbial communities for butyrate production to illustrate the use of inhomogeneous constraints to specify design problems. Starting from a community of 25 microbes, MPs enabled complete enumeration of communities for a range of minimal butyrate production requirements. Note that, in contrast to statistical models, good agreement with experimental butyrate production data did not require training or parameter estimation from such data, except for a rough estimate of minimal growth requirements.

Overall, we argue that MPs open up new possibilities for the detailed analysis of large-scale metabolic networks. Given the biological realism of MPs demonstrated in our example applications, we envisage MPs as particularly useful for the generation of experimentally testable hypotheses. For example, MP-based predictions could guide the design of targeted *in vitro* or *in vivo* experiments to elucidate metabolite exchanges in microbial communities. In addition—and less mechanistically—one could use MPs to design informative experiments for the learning of statistical models. Directly or indirectly, we thereby expect MPs to produce new biological insights into uni- and multicellular systems.

## Methods

### Constraint-based models and flux balance analysis

We consider constraint-based models (CBMs) at steady state described by a linear system of *m* metabolite mass balances with *n* fluxes **r** ∈ ℝ^*n*^ constrained by the stoichiometric matrix **N** ∈ ℝ^*m*× *n*^,

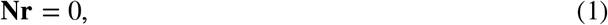

and any other linear constraints of the form

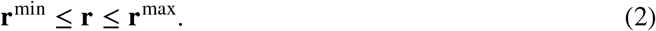

Flux balance analysis (FBA)^14^ maximizes or minimizes an assumed objective subject to constraints (1) and (2) in a linear program (LP):

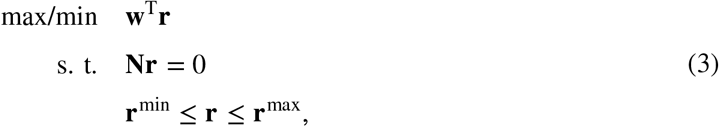

where **w** is the vector of weights for the fluxes in the objective.

### Minimal pathways and cut sets

We define the metabolic network *𝒩* = {1, 2, …, *n*} as the set of reaction indices in **r**, and we define a subnetwork *𝒮* and a pathway 𝒫 such that 𝒫 ⊆ 𝒮 ⊆ *𝒩*. We further require 𝒫 to be a minimal pathway (MP), here defined as a minimal set of reactions in 𝒮 that need to be active (have non-zero flux) in order to satisfy (1), (2), and any other linear constraints on 𝒩. In other words, 𝒫 is support-minimal: if the only active reactions in 𝒮 are 𝒫, constraints on 𝒩 will be violated if any reaction in 𝒫 is deactivated. We denote the set of known MPs as Π.

We define a cut set 𝒞 ⊆ ⋃ Π as a set of reactions that intersects every 𝒫 ∈ Π. This means that deactivating all reactions in 𝒞 will deactivate all known MPs. We also require 𝒞 to be support-minimal such that removing any reaction from 𝒞 will leave an empty intersection with at least one 𝒫 ∈ Π. A minimal cut set (MCS) deactivates all MPs in 𝒮 and thereby disables flux in 𝒩 as a whole. We denote the set of known MCSs as Γ.

### Model preprocessing

We use flux variability analysis (FVA)^15^ to determine tight bounds 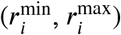 on feasible fluxes for all *i* ∈ 𝒩. Reactions in *𝒮* are then made irreversible with positive upper bounds by replacing reversible reactions by two irreversible ones and reversing irreversible reactions with negative lower bounds. FVA also reveals blocked reactions,

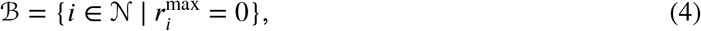

which always have zero flux and therefore are not part of any MP or MCS, and essential reactions,

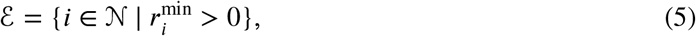

which cannot have zero flux and therefore are MCSs that participate in all MPs. Hence, only the subnetwork 𝒮 \ (ℬ ∪ℰ) can form MPs and MCSs along with ℰ. For simplicity, we will keep denoting this subnetwork as 𝒮.

### Finding a new pathway with direct minimization

For each *i* ∈ *𝒩*, we use a binary variable *b*_*i*_ ∈ {0, 1} to indicate if **r**_*i*_ is non-zero, as ensured by an inequality constraint:

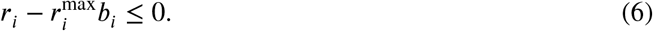

Another inequality constraint stops each 𝒫 ∈ Π from being found again:

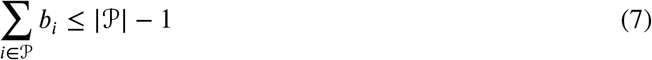

We can combine constraints (2), (6), and (7):

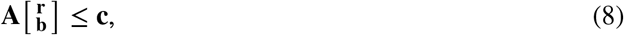

where the matrix **A** contains the coefficients, **b** is the vector of indicator variables, and **c** is the vector of constant terms. A new MP is then found by solving the MILP

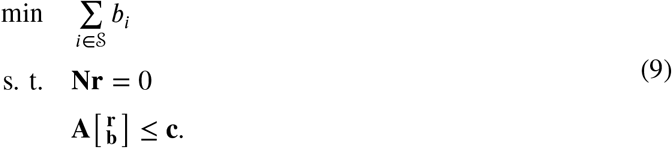

It minimizes the number of active reactions in 𝒮 by minimizing the number of non-zero indicator variables. In an optimal solution, the active reactions in 𝒮 define an MP that has the smallest possible number of active reactions among all MPs that have not yet been found.

### Finding a new pathway with iterative minimization

Assuming that a cut set that disables known MPs has been applied, our iterative method (**Algorithm S1**) does not require integer variables or additional constraints to find a new MP. At each iteration, a reaction *i* ∈ 𝒮 is deactivated before minimizing the sum of fluxes in 𝒮 in the LP

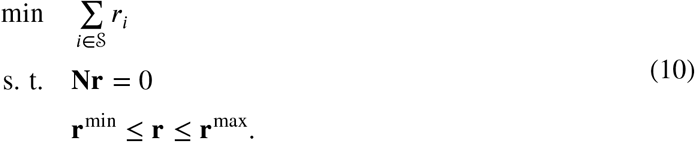

If an optimal solution is found, reactions in 𝒮 with zero flux in the solution can be deactivated as well because they are not needed in 𝒫. If no optimal solution is found, *i* must be part of 𝒫 and is reactivated. We repeat this until all *i* ∈ 𝒮have been tested, and the remaining active reactions in 𝒮 constitute an MP that is support-minimal but does not necessarily have the smallest possible number of reactions.

We can also randomize the order of MPs by deactivating a random *i* ∈ 𝒮 at each iteration. In this case, we cannot deactivate all reactions in 𝒮 with zero flux in an optimal solution because randomness would not be preserved. However, random reactions can be drawn and deactivated until a reaction with non-zero flux in the optimal solution is drawn.

### Finding a new cut set

We find a new cut set that deactivates all known MPs as described for EFMs by Song et al.^45^. Each reaction in at least one known MP, *i* ∈ ∪ Π, is represented by a binary variable *b*_*i*_ ∈ {0, 1}, and we use a constraint for each 𝒫 ∈ Π to ensure that it is covered by the cut set:

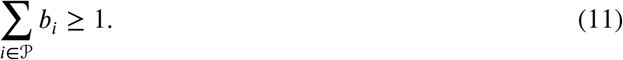

We also use a constraint for each 𝒞 ∈ Γ to exclude known MCSs:

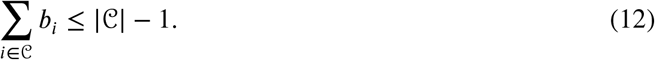

By reformulating constraints (11) and (12) as in (8), a new cut set results from solving the binary integer program (BIP)

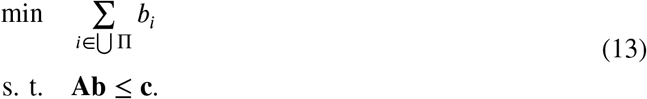

Specifically, a new cut set is given by the non-zero binary variables in the optimal solution; it may or may not be an MCS.

To find a new MP with iterative minimization, it is necessary to find a *viable* cut set that intersects all 𝒫 ∈ Π but does not intersect at least one unknown MP (i.e., a cut set that is not an MCS). To achieve this, **Algorithm S2** repeatedly solves and constrains the BIP and checks viability with the LP.

### Simple enumeration and sampling of pathways

Direct minimization can be used to enumerate MPs in order of increasing size by repeating the following steps until the MILP becomes infeasible:

1. Find a new MP by solving the MILP.
2. Add constraint (7) to the MILP to avoid finding the MP again.

Iterative minimization can be combined with the BIP for finding new cut sets to enumerate MPs based on the procedure by Song et al.^45^. The following steps are repeated until the BIP becomes infeasible:

1. Find and apply a new viable cut set using **Algorithm S2**.
2. Find a new MP using **Algorithm S1**.
3. Add constraint (11) to the BIP to avoid finding the cut set again.

This produces all MPs (not in order of increasing size) as well as all MCSs.

If too many MPs exist for complete enumeration to be feasible, MPs can be sampled randomly by repeatedly finding random MPs without finding and applying cut sets until the number of MPs equals the desired sample size. This avoids the significant overhead of solving BIPs but risks finding the same MP multiple times if the number of MPs is not sufficiently large.

### Graph-based enumeration of pathways

An undirected graph can be used to predict new MPs from known MPs. We define this graph as the union of reaction pairs from all 𝒫 ∈ Π, 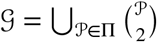, where reactions are nodes and reaction pairs are edges connecting two nodes. Reactions are connected if they occur together in at least one known MP, meaning that MPs are *cliques* (complete subgraphs) in 𝒢. We exploit that edges defined by known MPs can combine with each other to form additional cliques that may contain previously unknown MPs.

Each time we find a new MP during enumeration, we update the graph by adding any missing edges between reactions in the new MP. We use a modified Bron-Kerbosch algorithm^81^ to find *maximal cliques*, i.e., fully connected subgraphs where no other nodes in the graph are connected to all nodes in the subgraph. The MPs in each new maximal clique are enumerated separately using iterative minimization with the clique as subnetwork in place of 𝒮. This reduces the sizes of the optimization problems and thereby accelerates enumeration.

Graph-based prediction of new MPs can be combined with direct or iterative minimization (or any other procedure that returns new MPs); here, we implemented iterative minimization as described in **Algorithm S3**. However, graph-based prediction is biased by the known MPs and therefore incompatible with random sampling of MPs.

### Comparing MPs and other pathway definitions

First, we determined the key properties of EFMs, EFVs, ECMs, ProCEMs, EFPs, and MRSs from published theory and compared them to our definition of MPs. For each pathway definition, we checked whether it (1) allowed inhomogeneous constraints, (2) allowed targeted analysis of subnetworks, (3) required support-minimality, and (4) included flux stoichiometries or only patterns. We also identified the state-of-the-art (i.e., the most efficient) existing methods for enumeration from literature. Second, we built a CBM of the example network in **Fig. 1** using COBRApy^82^ and exported it to MATLAB format. We used CellNetAnalyzer^83^ (v2023.1) to load the model in MATLAB and enumerate the EFMs and EFVs of the example network. Specifically, we used the “CNAcobra2cna” function to convert the model to CellNetAnalyzer format and the “CNAcomputeEFM” function to enumerate EFMs and EFVs, using only binary reversibility constraints for EFM enumeration while specifying flux bounds for EFV enumeration. Finally, to compare scaling of pathway counts with network and subnetwork size, we collected data from complete enumerations of EFMs, EFVs, ECMs, EFPs, and MRSs reported in published studies. We did this manually by searching Google Scholar and extracting network size, subnetwork size in the case of ECMs and EFPs, and number of enumerated pathways.

### Benchmarking

We obtained six CBMs from the BiGG database^10^ (**Table S1**), comprising models of *Escherichia coli*^46,84^, *Helicobacter pylori*^85^, *Saccharomyces cerevisiae*^86^, *Cricetulus griseus*^87^, *and Homo sapiens*^59^. We removed any non-growth-associated maintenance reactions and all biomass reactions except the one with the largest number of metabolites, set the default bounds of fluxes to ± 1,000 mmol gDW^−1^ h^−1^, set a minimal growth rate of 0.1 h^−1^, and allowed uptake and secretion of all environmental metabolites to simulate a complex medium. We preprocessed each model as described above. For each model, we sampled random subnetworks ranging from ten reactions to the full network. We chose logarithmically spaced subnetwork sizes with four sizes sampled for each order of magnitude between 10 and 10^4^. We sampled 100 random subnetworks of each size and enumerated MPs using direct and iterative minimization (with and without graph or randomization). Incomplete enumerations were stopped after one hour. We used bootstrapping to compute 95% confidence intervals for estimated means. The data were resampled 1,000 times with replacement, the mean was reestimated for each sample, and a 95% confidence interval was defined from the 2.5th and 97.5th percentiles of these estimates. We compared means using a t-test or one-way ANOVA, and we used the Benjamini-Hochberg method to adjust for multiple testing^88^.

### *Application to E. coli* central carbon metabolism

We obtained a pathway map of *E. coli* central carbon metabolism from Escher^54^ and used the reactions in this map as a subnetwork within the genome-scale model iJO1366^53^. We also obtained single gene knockout data for a minimal glucose medium from Baba et al.^48^ and multiple gene knockout data for a non-minimal glucose and amino acid medium from Nakahigashi et al.^49^. Using randomized iterative minimization, we sampled 100,000 MPs in the subnetwork for each medium with a growth requirement of 0.1 h^−1^ and aerobic conditions. We used the default minimal glucose medium in the model for comparison to the single knockout data and the reported glucose and amino acid medium for comparison to the multiple knockout data^49^. We defined reaction and gene frequencies as the fractions of sampled MPs requiring each reaction or enzyme-encoding gene, respectively. To convert samples from reactions to the gene sets tested in knockout experiments, we computed the contribution of each gene set to each reaction by dividing the number of isozymes requiring at least one gene in the set by the total number of isozymes catalyzing the reaction. The contribution of each gene set to each sample was then computed as the maximum fraction of isozymes requiring at least one gene in the set across reactions in the sample. We also predicted the effects of single and multiple gene knockouts with MOMA^56^, using the “single_gene_deletion” function in COBRApy^82^. To compare MOMA predictions to gene frequencies from sampled MPs, we divided growth rates predicted by MOMA by the wild-type growth rate predicted by FBA. Bootstrapping was performed as for benchmarking.

### Application to host-microbe metabolite exchanges in the human gut

We built a host-microbe model in which genome-scale CBMs of a human cell and of six commensal gut microbes (*Bacteroides thetaiotaomicron, E. coli, Faecalibacterium prausnitzii, Lactobacillus plantarum, Lactococcus lactis*, and *Streptococcus thermophilus*; **Table S2**) can exchange metabolites through the gut lumen. We based the model on previous work^57^ but used a more recent human model^59^ and collection of gut microbe models^58^, all obtained from Virtual Metabolic Human^89^. We set a non-growth-associated ATP requirement of 0.1 mmol gDW^−1^ h^−1^ for the human cell and a growth requirement of 0.1 h^−1^ for the microbes. To simulate the anaerobic and potentially complex metabolic environment of the gut, we allowed uptake and secretion of all environmental metabolites for all cells, turned off oxygen exchange for the microbes, and added reactions representing all possible metabolite exchanges between microbes and from microbes to the human cell (excluding ATP, sugars, and inorganic ions and compounds). Building on our recent approach^37^, we also applied two FBA-steps, minimizing (1) microbial intracellular flux^60^ and (2) uptake flux not explained by metabolite exchange across all cells:

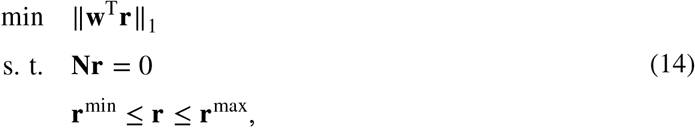

where ∥ **w**^T^**r**∥_1_ is the weighted sum of absolute fluxes to be minimized. After the first step, we constrained each intracellular microbial flux to its value in the minimum obtained. After the second step, we constrained the absolute sum of exchange fluxes not explained by metabolite exchange to the minimum obtained, 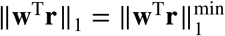. This approach aimed to explain as much of the functional requirements as possible in terms of metabolite cross-feeding with biologically reasonable microbial flux distributions. We enumerated minimal sets of metabolite exchanges using iterative minimization with graph.

### Application to butyrate-producing microbial communities

We built a community model using genome-scale CBMs of the 25 gut microbes cultivated by Clark et al.^73^ from the AGORA collection^58^ (**Table S3**). An *in silico* butyrate production medium, similar to the *in vitro* medium used by Clark et al.^73^, was created. As a starting point, we used the *western diet* medium^57^, in which all AGORA models are known to grow anaerobically^58^. Essential compounds such as vitamins and minerals were left untouched, whereas the supply of sugars, fibers, lipids, and amino acids were adapted to the experimental concentrations and normalized, such that the summed supply of amino acids in the western diet equals the sum of amino acids in the constructed medium. The 25 models were constrained to the butyrate production medium by restricting the extracellular input fluxes to be smaller or equal to the given medium constraints. In a few cases, compounds outside the butyrate production medium were supplied in small quantities to allow for simulated growth.

Lactate metabolism was curated for the 25 strains. Several strain models included lactose dehydrogenase (LDH) with equation lactate + NAD^+^ ↔pyruvate + NADH. **Table S4** shows the reaction energy of LDH, computed with eQuilibrator^90^ for standard conditions and for the smallest ratio of lactate to pyruvate that renders the reaction feasible. For the feasibility calculation, measured NAD^+^ and NADH concentrations of *E. coli* were used^91^. The default eQuilibrator parameters pH 7.5, pMg 3, and ionic strength 0.25 were used^90^. The calculations show that, to reach Δ_**r**_*G*^′^ = 0, the intracellular lactate concentration would have to be three orders of magnitude higher than the pyruvate concentration, which is deemed unlikely. Due to this, wherever LDH occured in the models, it was restricted to only operate in the backward direction. *Anaerostipes caccae* is a known lactose consumer. Following KEGG annotation for genome T06754, *Anaerostipes caccae* encodes enzyme EC 1.1.1.436, which is a lactate dehydrogenase with ferredoxin coupling, lactate+2NAD^+^ +ferredoxin^2–^ ↔ pyruvate+2 NADH+ferredoxin, known to operate in the forward direction^92^.

We were interested in finding minimal communities with high butyrate production capabilites. Since the primary interest was not in *how* this is achieved intracellularly, we simplified the interiors of the cells in a linear fashion to reduce the number of reactions and thereby speed up calculations. In general, due to constraint (1), the linear number of degrees of freedom of **r** is *n*_*u*_ = *n* − *rank*(**N**). By computing the reduced row echelon form (RREF) of **N**, an unconstrained basis 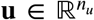 is attained, reducing the number of fluxes by *rank*(**N**), which is typically more than half of the fluxes. RREF, rather than SVD, is used so that the transformation matrix **Tr = u** remains sparse. To ensure numeric stability of the RREF computation, it was performed symbolically on a slightly altered form of **N**, in which all elements are truncated to have no more than two digits. The basis **u** represents a subset of the fluxes **r**. Since some of the fluxes of interest, in this case exchange fluxes, may be absent in **u**, they were added to **u** in a post-processing step and constrained to be identical to their corresponding reactions in **r**.

We required all strains participating in a sub-community to grow efficiently by constraining the growth rate to be larger than a fraction *α* ∈ [0, 1] of the sum of absolute fluxes (in *u* space). To validate the genome-scale modeling approach in the butyrate production setting and to estimate *α*, we used 1,799 out of 1,850 *in vitro* experiments with different community compositions performed by Clark et al.^73^. The 51 experiments that were not considered either had insufficient reads for relative species abundance quantification or more than 1% of the relative abundance was occupied by a species outside the catalog of 25 species. For model validation, qualitative community composition was used, with a threshold of 1% relative abundance required for a species to count as included in an experiment. After applying the threshold, 790 qualitatively distinct communities remained and were assigned the average butyrate production of the participating experiments. Maximal butyrate production was simulated using FBA for each of the experimentally tested communities. The highest correlation between simulated butyrate flux and measured butyrate concentration was 0.59 at *α* = 0.8. Weighted Pearson correlation was used, with weights equal to the number of butyrate measurements underlying average butyrate production for the 790 qualitatively distinct communities.

We performed community enumeration using iterative minimization with graph, *α* = 0.8, and the set of biomass reactions as subnetwork. We required butyrate production of 50%, 75%, 90%, and 95% of the maximal butyrate production rate of the full 25-strain community. To assess significance of the enumerated minimal butyrate-producing communities, we compared them to randomized communities. Specifically, we resampled 500 of the enumerated communities with replacement for each level. For each community, we kept the butyrate producers, replaced all other strains by random, non-butyrate-producing strains sampled without replacement, and maximized butyrate production.

## Data and code availability

Enumeration and sampling of MPs was implemented in the Python package *mptool*, which is available at https://gitlab.com/csb.ethz/mptool along with data and code needed to reproduce our results.

## Acknowledgements

This work was supported by the SystemsX.ch project GutX, evaluated by the Swiss National Science Foundation, and the Swiss National Science Foundation Sinergia project #177164. We thank Ylva Katarina Wedmark, Jon Olav Vik, Filip Rotnes, Mattia Gollub, and Constance Le Gac for helpful comments on the manuscript.

## Author contributions

J.S. and O.Ø. designed the study. O.Ø. developed the algorithms. O.Ø. and A.T. performed analyses. O.Ø., A.T., and J.S. wrote the manuscript. All authors read and approved the final manuscript.

## References

[1] Skwara, A., Gowda, K., Yousef, M., Diaz-Colunga, J., Raman, A.S., Sanchez, A., Tikhonov, M., and Kuehn, S. Statistically learning the functional landscape of microbial communities. Nature Ecology and Evolution, 7(11):1823–1833, 2023.

[2] Arya, S., George, A.B., and O’Dwyer, J.P. Sparsity of higher-order landscape interactions enables learning and prediction for microbiomes. Proceedings of the National Academy of Sciences, 120 (48), 2023.

[3] Diaz-Colunga, J., Skwara, A., Vila, J.C.C., Bajic, D., and Sanchez, A. Global epistasis and the emergence of function in microbial consortia. Cell, 187(12):3108–3119.e30, 2024.

[4] Visconti, A., Le Roy, C.I., Rosa, F., Rossi, N., Martin, T.C., Mohney, R.P., Li, W., de Rinaldis, E., Bell, J.T., Venter, J.C., Nelson, K.E., Spector, T.D., and Falchi, M. Interplay between the human gut microbiome and host metabolism. Nature Communications, 10(1), 2019.

[5] Yilmaz, B., Juillerat, P., Øyås, O., Ramon, C., Bravo, F.D., Franc, Y., Fournier, N., Michetti, P., Mueller, C., Geuking, M., Pittet, V.E.H., Maillard, M.H., Rogler, G., Swiss IBD Cohort Investigators, Wiest, R., Stelling, J., and Macpherson, A.J. Microbial network disturbances in relapsing refractory Crohn’s disease. Nature Medicine, 25(2), 2019.

[6] Estrela, S., Vila, J.C.C., Lu, N., Baji, D., Rebolleda-Gómez, M., Chang, C.Y., Goldford, J.E., Sanchez-Gorostiaga, A., and Álvaro Sánchez. Functional attractors in microbial community assembly. Cell Systems, 13(1):29–42.e7, 2022.

[7] Chang, C.Y., Baji, D., Vila, J.C.C., Estrela, S., and Sanchez, A. Emergent coexistence in multispecies microbial communities. Science, 381(6655):343–348, 2023.

[8] Pontrelli, S., Szabo, R., Pollak, S., Schwartzman, J., Ledezma-Tejeida, D., Cordero, O.X., and Sauer, U. Metabolic cross-feeding structures the assembly of polysaccharide degrading communities. Science Advances, 8(8), 2022.

[9] Ganter, M., Bernard, T., Moretti, S., Stelling, J., and Pagni, M. MetaNetX.org: a website and repository for accessing, analysing and manipulating metabolic networks. Bioinformatics, 29(6), 2013.

[10] Norsigian, C.J., Pusarla, N., McConn, J.L., Yurkovich, J.T., Dräger, A., Palsson, B.O., and King, Z. BiGG Models 2020: Multi-strain genome-scale models and expansion across the phylogenetic tree. Nucleic Acids Research, 48(D1), 2020.

[11] Varma, A. and Palsson, B.O. Metabolic flux balancing: Basic concepts, scientific and practical use. Bio/Technology, 12(10), 1994.

[12] Urbanczik, R. Enumerating constrained elementary flux vectors of metabolic networks. IET Systems Biology, 1(5), 2007.

[13] Klamt, S., Regensburger, G., Gerstl, M.P., Jungreuthmayer, C., Schuster, S., Mahadevan, R., Zanghellini, J., and Müller, S. From elementary flux modes to elementary flux vectors: Metabolic pathway analysis with arbitrary linear flux constraints. PLOS Computational Biology, 13(4), 2017.

[14] Orth, J.D., Thiele, I., and Palsson, B.O. What is flux balance analysis? Nature Biotechnology, 28 (3), 2010.

[15] Mahadevan, R. and Schilling, C.H. The e”ects of alternate optimal solutions in constraint-based genome-scale metabolic models. Metabolic Engineering, 5(4), 2003.

[16] Schuster, S. and Hilgetag, C. On elementary flux modes in biochemical systems at steady state. Journal of Biological Systems, 2(2), 1994.

[17] Klamt, S. and Stelling, J. Combinatorial complexity of pathway analysis in metabolic networks. Molecular Biology Reports, 29(1-2), 2002.

[18] Acuña, V., Chierichetti, F., Lacroix, V., Marchetti-Spaccamela, A., Sagot, M.F., and Stougie, L. Modes and cuts in metabolic networks: Complexity and algorithms. BioSystems, 95(1), 2009.

[19] Ullah, E., Yosafshahi, M., and Hassoun, S. Towards scaling elementary flux mode computation. Briefings in Bioinformatics, 21(6), 2020.

[20] Terzer, M. and Stelling, J. Large-scale computation of elementary flux modes with bit pattern trees. Bioinformatics, 24(19), 2008.

[21] Hunt, K.A., Folsom, J.P., Ta”s, R.L., and Carlson, R.P. Complete enumeration of elementary flux modes through scalable demand-based subnetwork definition. Bioinformatics, 30(11), 2014.

[22] van Klinken, J.B. and van Dijk, K.W. FluxModeCalculator: An e!cient tool for large-scale flux mode computation. Bioinformatics, 32(8), 2016.

[23] Buchner, B.A. and Zanghellini, J. EFMlrs: a Python package for elementary flux mode enumeration via lexicographic reverse search. BMC Bioinformatics, 22(1), 2021.

[24] Zanghellini, J., Gerstl, M.P., Hanscho, M., Nair, G., Regensburger, G., Müller, S., and Jungreuthmayer, C. Toward Genome-Scale Metabolic Pathway Analysis. In Industrial Biotechnology: Microorganisms. 2016.

[25] de Figueiredo, L.F., Podhorski, A., Rubio, A., Kaleta, C., Beasley, J.E., Schuster, S., and Planes, F.J. Computing the shortest elementary flux modes in genome-scale metabolic networks. Bioinformatics, 25(23), 2009.

[26] Machado, D., Soons, Z., Patil, K.R., Ferreira, E.C., and Rocha, I. Random sampling of elementary flux modes in large-scale metabolic networks. Bioinformatics, 28(18), 2012.

[27] David, L. and Bockmayr, A. Computing elementary flux modes involving a set of target reactions. IEEE/ACM Transactions on Computational Biology and Bioinformatics, 11(6), 2014.

[28] Rezola, A., de Figueiredo, L.F., Brock, M., Pey, J., Podhorski, A., Wittmann, C., Schuster, S., Bockmayr, A., and Planes, F.J. Exploring metabolic pathways in genome-scale networks via generating flux modes. Bioinformatics, 27(4), 2011.

[29] Röhl, A. and Bockmayr, A. Finding MEMo: minimum sets of elementary flux modes. Journal of Mathematical Biology, 79(5), 2019.

[30] Jol, S.J., Kümmel, A., Terzer, M., Stelling, J., and Heinemann, M. System-level insights into yeast metabolism by thermodynamic analysis of elementary flux modes. PLOS Computational Biology, 8(3), 2012.

[31] Jungreuthmayer, C., Ruckerbauer, D.E., and Zanghellini, J. RegEfmtool: Speeding up elementary flux mode calculation using transcriptional regulatory rules in the form of three-state logic. BioSystems, 113(1), 2013.

[32] Gerstl, M.P., Ruckerbauer, D.E., Mattanovich, D., Jungreuthmayer, C., and Zanghellini, J. Metabolomics integrated elementary flux mode analysis in large metabolic networks. Scientific Reports, 5(8930), 2015.

[33] Marashi, S.A., David, L., and Bockmayr, A. Analysis of Metabolic Subnetworks by Flux Cone Projection. Algorithms for Molecular Biology, 7(1), 2012.

[34] Urbanczik, R. and Wagner, C. Functional stoichiometric analysis of metabolic networks. Bioinformatics, 21(22), 2005.

[35] Clement, T.J., Baalhuis, E.B., Teusink, B., Bruggeman, F.J., Planqué, R., and de Groot, D.H. Unlocking Elementary Conversion Modes: ecmtool Unveils All Capabilities of Metabolic Networks. Patterns, 2(1), 2021.

[36] Kaleta, C., De Figueiredo, L.F., and Schuster, S. Can the whole be less than the sum of its parts? Pathway analysis in genome-scale metabolic networks using elementary flux patterns. Genome Research, 19(10), 2009.

[37] Øyås, O., Borrell, S., Trauner, A., Zimmermann, M., Feldmann, J., Liphardt, T., Gagneux, S., Stelling, J., Sauer, U., and Zampieri, M. Model-based integration of genomics and metabolomics reveals SNP functionality in Mycobacterium tuberculosis. Proceedings of the National Academy of Sciences, 117(15), 2020.

[38] Burgard, A.P., Vaidyaraman, S., and Maranas, C.D. Minimal reaction sets for Escherichia coli metabolism under di”erent growth requirements and uptake environments. Biotechnology Progress, 17(5), 2001.

[39] Jonnalagadda, S. and Srinivasan, R. An efficient graph theory based method to identify every minimal reaction set in a metabolic network. BMC Systems Biology, 8(1), 2014.

[40] Röhl, A. and Bockmayr, A. A mixed-integer linear programming approach to the reduction of genome-scale metabolic networks. BMC Bioinformatics, 18(1):1–10, 2017.

[41] Pey, J. and Planes, F.J. Direct calculation of elementary flux modes satisfying several biological constraints in genome-scale metabolic networks. Bioinformatics, 30(15), 2014.

[42] Quek, L.E. and Nielsen, L.K. A depth-first search algorithm to compute elementary flux modes by linear programming. BMC Systems Biology, 8(1), 2014.

[43] Pey, J., Villar, J.A., Tobalina, L., Rezola, A., García, J.M., Beasley, J.E., and Planes, F.J. TreeEFM: calculating elementary flux modes using linear optimization in a tree-based algorithm. Bioinformatics, 31(6):897–904, 2015.

[44] Buchner, B., Clement, T.J., de Groot, D.H., and Zanghellini, J. ecmtool: fast and memory-e!cient enumeration of elementary conversion modes. Bioinformatics, 39(3):btad095, 2023.

[45] Song, H.S., Goldberg, N., Mahajan, A., and Ramkrishna, D. Sequential computation of elementary modes and minimal cut sets in genome-scale metabolic networks using alternate integer linear programming. Bioinformatics, 33(15), 2017.

[46] Monk, J.M., Lloyd, C.J., Brunk, E., Mih, N., Sastry, A., King, Z., Takeuchi, R., Nomura, W., Zhang, Z., Mori, H., Feist, A.M., and Palsson, B.O. iML1515, a knowledgebase that computes Escherichia coli traits. Nature Biotechnology, 35(10), 2017.

[47] Rancati, G., Mo”at, J., Typas, A., and Pavelka, N. Emerging and evolving concepts in gene essentiality. Nature Reviews Genetics, 19(1), 2018.

[48] Baba, T., Ara, T., Hasegawa, M., Takai, Y., Okumura, Y., Baba, M., Datsenko, K.A., Tomita, M., Wanner, B.L., and Mori, H. Construction of Escherichia coli K-12 in-frame, single-gene knockout mutants: The Keio collection. Molecular Systems Biology, 2(1), 2006.

[49] Nakahigashi, K., Toya, Y., Ishii, N., Soga, T., Hasegawa, M., Watanabe, H., Takai, Y., Honma, M., Mori, H., and Tomita, M. Systematic phenome analysis of escherichia coli multiple-knockout mutants reveals hidden reactions in central carbon metabolism. Molecular Systems Biology, 5(1): 306, 2009.

[50] Goodall, E.C., Robinson, A., Johnston, I.G., Jabbari, S., Turner, K.A., Cunningham, A.F., Lund, P.A., Cole, J.A., and Henderson, I.R. The essential genome of Escherichia coli K-12. mBio, 9(1), 2018.

[51] Wang, T., Guan, C., Guo, J., Liu, B., Wu, Y., Xie, Z., Zhang, C., and Xing, X.H. Pooled CRISPR interference screening enables genome-scale functional genomics study in bacteria with superior performance. Nature Communications, 9(1), 2018.

[52] Rousset, F., Cui, L., Siouve, E., Becavin, C., Depardieu, F., and Bikard, D. Genome-wide CRISPR-dCas9 screens in E. coli identify essential genes and phage host factors. PLOS Genetics, 14(11), 2018.

[53] Orth, J.D., Conrad, T.M., Na, J., Lerman, J.A., Nam, H., Feist, A.M., and Palsson, B. A comprehensive genome-scale reconstruction of Escherichia coli metabolism-2011. Molecular Systems Biology, 7(535), 2011.

[54] King, Z.A., Dräger, A., Ebrahim, A., Sonnenschein, N., Lewis, N.E., and Palsson, B.O. Escher: A Web Application for Building, Sharing, and Embedding Data-Rich Visualizations of Biological Pathways. PLOS Computational Biology, 11(8), 2015.

[55] Fischer, E. and Sauer, U. A novel metabolic cycle catalyzes glucose oxidation and anaplerosis in hungry Escherichia coli. The Journal of Biological Chemistry, 278(47), 2003.

[56] Segre, D., Vitkup, D., and Church, G.M. Analysis of optimality in natural and perturbed metabolic networks. Proceedings of the National Academy of Sciences, 99(23):15112–15117, 2002.

[57] Heinken, A. and Thiele, I. Systematic prediction of health-relevant human-microbial cometabolism through a computational framework. Gut Microbes, 6(2), 2015.

[58] Magnúsdóttir, S., Heinken, A., Kutt, L., Ravcheev, D.A., Bauer, E., Noronha, A., Greenhalgh, K., Jäger, C., Baginska, J., Wilmes, P., Fleming, R.M., and Thiele, I. Generation of genome-scale metabolic reconstructions for 773 members of the human gut microbiota. Nature Biotechnology, 35(1), 2017.

[59] Brunk, E., Sahoo, S., Zielinski, D.C., Altunkaya, A., Dräger, A., Mih, N., Gatto, F., Nilsson, A., Preciat Gonzalez, G.A., Aurich, M.K., Prlic, A., Sastry, A., Danielsdottir, A.D., Heinken, A., Noronha, A., Rose, P.W., Burley, S.K., Fleming, R.M., Nielsen, J., Thiele, I., and Palsson, B.O. Recon3D enables a three-dimensional view of gene variation in human metabolism. Nature Biotechnology, 36(3), 2018.

[60] Lewis, N.E., Hixson, K.K., Conrad, T.M., Lerman, J.A., Charusanti, P., Polpitiya, A.D., Adkins, J.N., Schramm, G., Purvine, S.O., Lopez-Ferrer, D., Weitz, K.K., Eils, R., König, R., Smith, R.D., and Palsson, B. Omic data from evolved E. coli are consistent with computed optimal growth from genome-scale models. Molecular Systems Biology, 6(390), 2010.

[61] Machado, D. and Herrgård, M. Systematic Evaluation of Methods for Integration of Transcriptomic Data into Constraint-Based Models of Metabolism. PLOS Computational Biology, 10(4), 2014.

[62] Kumar, M., Ji, B., Babaei, P., Das, P., Lappa, D., Ramakrishnan, G., Fox, T.E., Haque, R., Petri, W.A., Bäckhed, F., and Nielsen, J. Gut microbiota dysbiosis is associated with malnutrition and reduced plasma amino acid levels: Lessons from genome-scale metabolic modeling. Metabolic Engineering, 49, 2018.

[63] Mardinoglu, A., Bergentall, M., Gha”ari, P., Larsson, E., Backhed, F., Shoaie, S., Nielsen, J., and Zhang, C. The gut microbiota modulates host amino acid and glutathione metabolism in mice. Molecular Systems Biology, 11(10), 2015.

[64] Zhu, L., Baker, S.S., Gill, C., Liu, W., Alkhouri, R., Baker, R.D., and Gill, S.R. Characterization of gut microbiomes in nonalcoholic steatohepatitis (NASH) patients: A connection between endogenous alcohol and NASH. Hepatology, 57(2), 2013.

[65] Abu-Shanab, A. and Quigley, E.M. The role of the gut microbiota in nonalcoholic fatty liver disease. Nature Reviews Gastroenterology & Hepatology, 7(12):691–701, 2010.

[66] De Weirdt, R., Possemiers, S., Vermeulen, G., Moerdijk-Poortvliet, T.C., Boschker, H.T., Verstraete, W., and Van de Wiele, T. Human faecal microbiota display variable patterns of glycerol metabolism. FEMS Microbiology Ecology, 74(3):601–611, 2010.

[67] Morrison, D.J. and Preston, T. Formation of short chain fatty acids by the gut microbiota and their impact on human metabolism. Gut Microbes, 7(3), 2016.

[68] Serena, C., Ceperuelo-Mallafré, V., Keiran, N., Queipo-Ortuño, M.I., Bernal, R., Gomez-Huelgas, R., Urpi-Sarda, M., Sabater, M., Pérez-Brocal, V., Andrés-Lacueva, C., Moya, A., Tinahones, F.J., Fernández-Real, J.M., Vendrell, J., and Fernández-Veledo, S. Elevated circulating levels of succinate in human obesity are linked to specific gut microbiota. ISME Journal, 12(7), 2018.

[69] Magnúsdóttir, S., Ravcheev, D., De Crécy-Lagard, V., and Thiele, I. Systematic genome assessment of B-vitamin biosynthesis suggests cooperation among gut microbes. Frontiers in Genetics, 6(148), 2015.

[70] Sharma, V., Rodionov, D.A., Leyn, S.A., Tran, D., Iablokov, S.N., Ding, H., Peterson, D.A., Osterman, A.L., and Peterson, S.N. B-Vitamin Sharing Promotes Stability of Gut Microbial Communities. Frontiers in Microbiology, 10(1485), 2019.

[71] Qi, H., Li, Y., Yun, H., Zhang, T., Huang, Y., Zhou, J., Yan, H., Wei, J., Liu, Y., Zhang, Z., Gao, Y., Che, Y., Su, X., Zhu, D., Zhang, Y., Zhong, J., and Yang, R. Lactobacillus maintains healthy gut mucosa by producing L-Ornithine. Communications Biology, 2(1), 2019.

[72] Louis, P. and Flint, H.J. Formation of propionate and butyrate by the human colonic microbiota. Environmental Microbiology, 19(1):29–41, 2017.

[73] Clark, R.L., Connors, B.M., Stevenson, D.M., Hromada, S.E., Hamilton, J.J., Amador-Noguez, D., and Venturelli, O.S. Design of synthetic human gut microbiome assembly and butyrate production. Nature Communications, 12(1):1–16, 2021.

[74] Duncan, S.H., Barcenilla, A., Stewart, C.S., Pryde, S.E., and Flint, H.J. Acetate utilization and butyryl coenzyme A (CoA): acetate-CoA transferase in butyrate-producing bacteria from the human large intestine. Applied and Environmental Microbiology, 68(10):5186–5190, 2002.

[75] Khan, M.T., Duncan, S.H., Stams, A.J., Van Dijl, J.M., Flint, H.J., and Harmsen, H.J. The gut anaerobe Faecalibacterium prausnitzii uses an extracellular electron shuttle to grow at oxic–anoxic interphases. The ISME journal, 6(8):1578–1585, 2012.

[76] Paczia, N., Nilgen, A., Lehmann, T., Gätgens, J., Wiechert, W., and Noack, S. Extensive exometabolome analysis reveals extended overflow metabolism in various microorganisms. Microbial cell factories, 11:1–14, 2012.

[77] Bishehsari, F., Magno, E., Swanson, G., Desai, V., Voigt, R.M., Forsyth, C.B., and Keshavarzian, A. Alcohol and gut-derived inflammation. Alcohol research: current reviews, 38(2):163, 2017.

[78] Seo, B., Jeon, K., Moon, S., Lee, K., Kim, W.K., Jeong, H., Cha, K.H., Lim, M.Y., Kang, W., Kweon, M.N., et al. Roseburia spp. abundance associates with alcohol consumption in humans and its administration ameliorates alcoholic fatty liver in mice. Cell host & microbe, 27(1):25–40, 2020.

[79] Schellenberger, J., Lewis, N.E., and Palsson, B. Elimination of thermodynamically infeasible loops in steady-state metabolic models. Biophysical Journal, 100(3), 2011.

[80] Ballerstein, K., von Kamp, A., Klamt, S., and Haus, U.U. Minimal cut sets in a metabolic network are elementary modes in a dual network. Bioinformatics, 28(3):381–387, 2012.

[81] Cazals, F. and Karande, C. A note on the problem of reporting maximal cliques. Theoretical Computer Science, 407(1-3), 2008.

[82] Ebrahim, A., Lerman, J.A., Palsson, B.O., and Hyduke, D.R. COBRApy: COnstraints-Based Reconstruction and Analysis for Python. BMC Systems Biology, 7(74), 2013.

[83] von Kamp, A., Thiele, S., Hädicke, O., and Klamt, S. Use of CellNetAnalyzer in biotechnology and metabolic engineering. Journal of Biotechnology, 261:221–228, 2017.

[84] Orth, J.D., Palsson, B.Ø., and Fleming, R.M.T. Reconstruction and Use of Microbial Metabolic Networks: the Core Escherichia coli Metabolic Model as an Educational Guide. EcoSal Plus, 4(1), 2010.

[85] Thiele, I., Vo, T.D., Price, N.D., and Palsson, B.Ø. Expanded metabolic reconstruction of Helicobacter pylori (iIT341 GSM/GPR): an in silico genome-scale characterization of single- and double-deletion mutants. Journal of Bacteriology, 187(16), 2005.

[86] Mo, M.L., Palsson, B.Ø., and Herrgård, M.J. Connecting extracellular metabolomic measurements to intracellular flux states in yeast. BMC Systems Biology, 3(37), 2009.

[87] Hefzi, H., Ang, K.S., Hanscho, M., Bordbar, A., Ruckerbauer, D., Lakshmanan, M., Orellana, C.A., Baycin-Hizal, D., Huang, Y., Ley, D., Martinez, V.S., Kyriakopoulos, S., Jiménez, N.E., Zielinski, D.C., Quek, L.E., Wul”, T., Arnsdorf, J., Li, S., Lee, J.S., Paglia, G., Loira, N., Spahn, P.N., Pedersen, L.E., Gutierrez, J.M., King, Z.A., Lund, A.M., Nagarajan, H., Thomas, A., Abdel-Haleem, A.M., Zanghellini, J., Kildegaard, H.F., Voldborg, B.G., Gerdtzen, Z.P., Betenbaugh, M.J., Palsson, B.O., Andersen, M.R., Nielsen, L.K., Borth, N., Lee, D.Y., and Lewis, N.E. A Consensus Genome-scale Reconstruction of Chinese Hamster Ovary Cell Metabolism. Cell Systems, 3(5), 2016.

[88] Benjamini, Y. and Hochberg, Y. Controlling the false discovery rate: a practical and powerful approach to multiple testing. Journal of the Royal Statistical Society: Series B (Methodological), 57(1):289–300, 1995.

[89] Noronha, A., Modamio, J., Jarosz, Y., Guerard, E., Sompairac, N., Preciat, G., Daníelsdóttir, A.D., Krecke, M., Merten, D., Haraldsdóttir, H.S., Heinken, A., Heirendt, L., Magnúsdóttir, S., Ravcheev, D.A., Sahoo, S., Gawron, P., Friscioni, L., Garcia, B., Prendergast, M., Puente, A., Rodrigues, M., Roy, A., Rouquaya, M., Wiltgen, L., $agare, A., John, E., Krueger, M., Kuperstein, I., Zinovyev, A., Schneider, R., Fleming, R.M., and Thiele, I. The Virtual Metabolic Human database: Integrating human and gut microbiome metabolism with nutrition and disease. Nucleic Acids Research, 47 (D1), 2019.

[90] Beber, M.E., Gollub, M.G., Moza”ari, D., Shebek, K.M., Flamholz, A.I., Milo, R., and Noor, E. eQuilibrator 3.0: a database solution for thermodynamic constant estimation. Nucleic Acids Research, 50(D1):D603–D609, 2022.

[91] Bennett, B.D., Kimball, E.H., Gao, M., Osterhout, R., Van Dien, S.J., and Rabinowitz, J.D. Absolute metabolite concentrations and implied enzyme active site occupancy in Escherichia coli. Nature Chemical Biology, 5(8):593–599, 2009.

[92] Wegho”, M.C., Bertsch, J., and Müller, V. A novel mode of lactate metabolism in strictly anaerobic bacteria. Environmental Microbiology, 17(3):670–677, 2015.

