## Supplementary Information for "Scalable enumeration and sampling of minimal metabolic pathways for organisms and communities"

### Supplementary figures

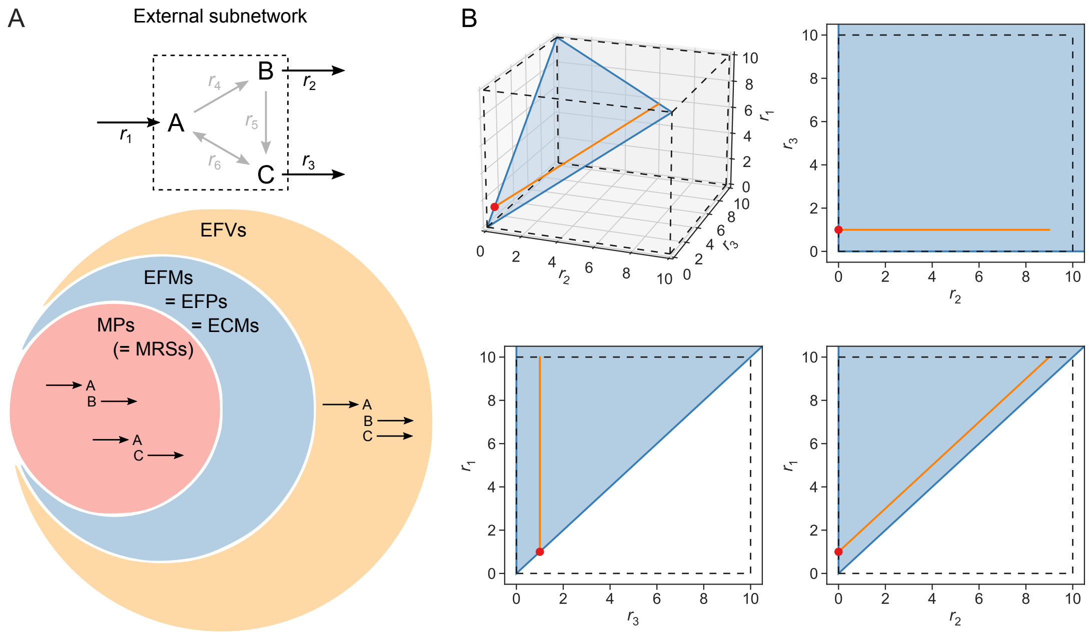


**Figure S1. Comparison of metabolic pathway definitions for an external subnetwork.** (**A**) Flux patterns for the subnetwork consisting of metabolites A–C and external reactions $r_1$–$r_3$ within the full network in **Fig. 1B**. (**B**) Three-dimensional flux space and its two-dimensional projections corresponding to EFMs (blue), EFVs (orange), and MPs (red) for the subnetwork from (A).

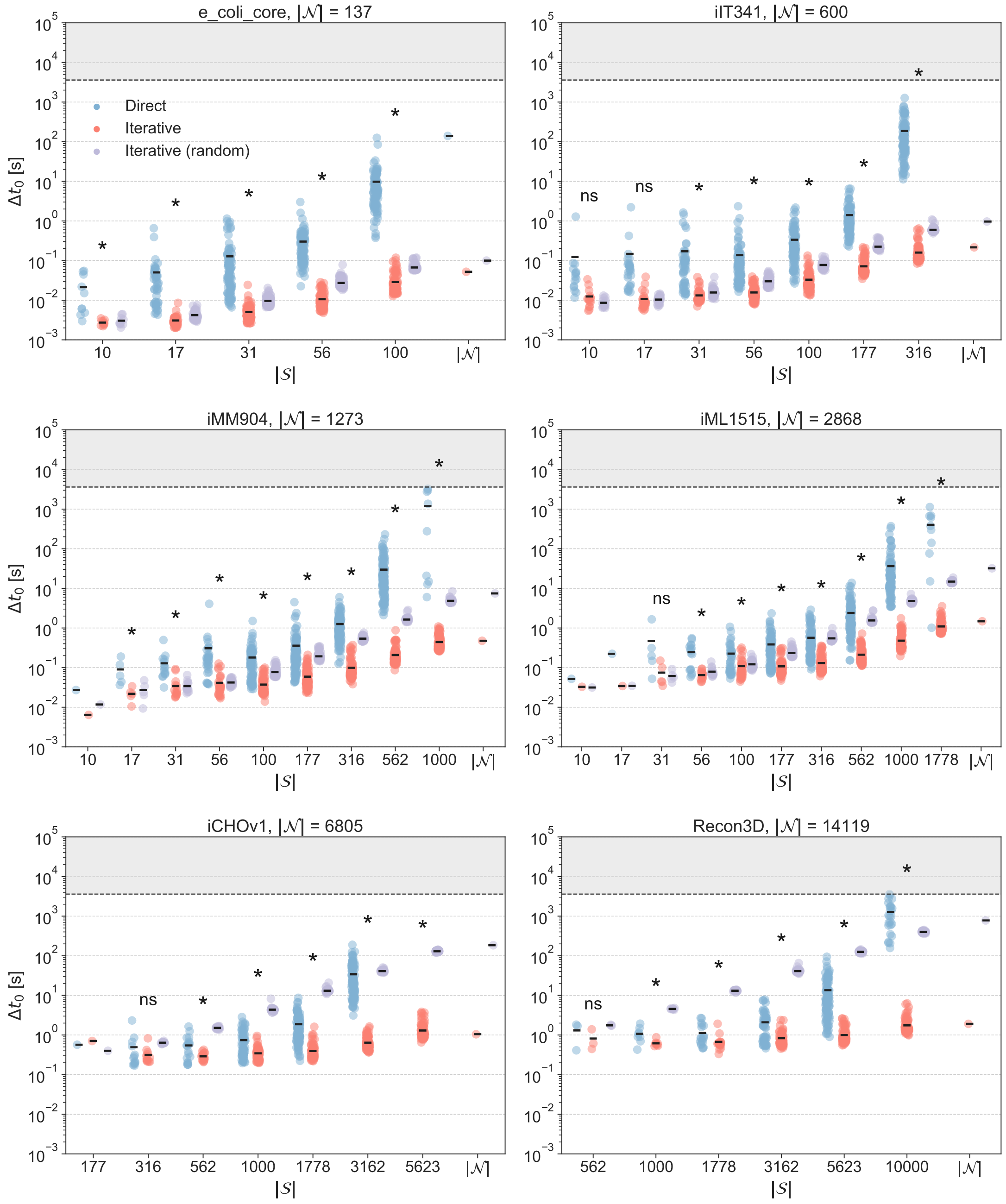


**Figure S2. Running time for finding the first MP.** Running time $\Delta t_0$ for finding the first new MP with direct (blue) and iterative minimization with (purple) and without (red) randomization for different subnetwork sizes $|\mathcal{S}|$ in six different models with network size $|\mathcal{N}|$. Stars indicate significant difference between means from one-way ANOVA where data was available for all three methods or $t$-test where data was available for two methods (significance level 0.05). Optimization was stopped after one hour (indicated by dashed line).

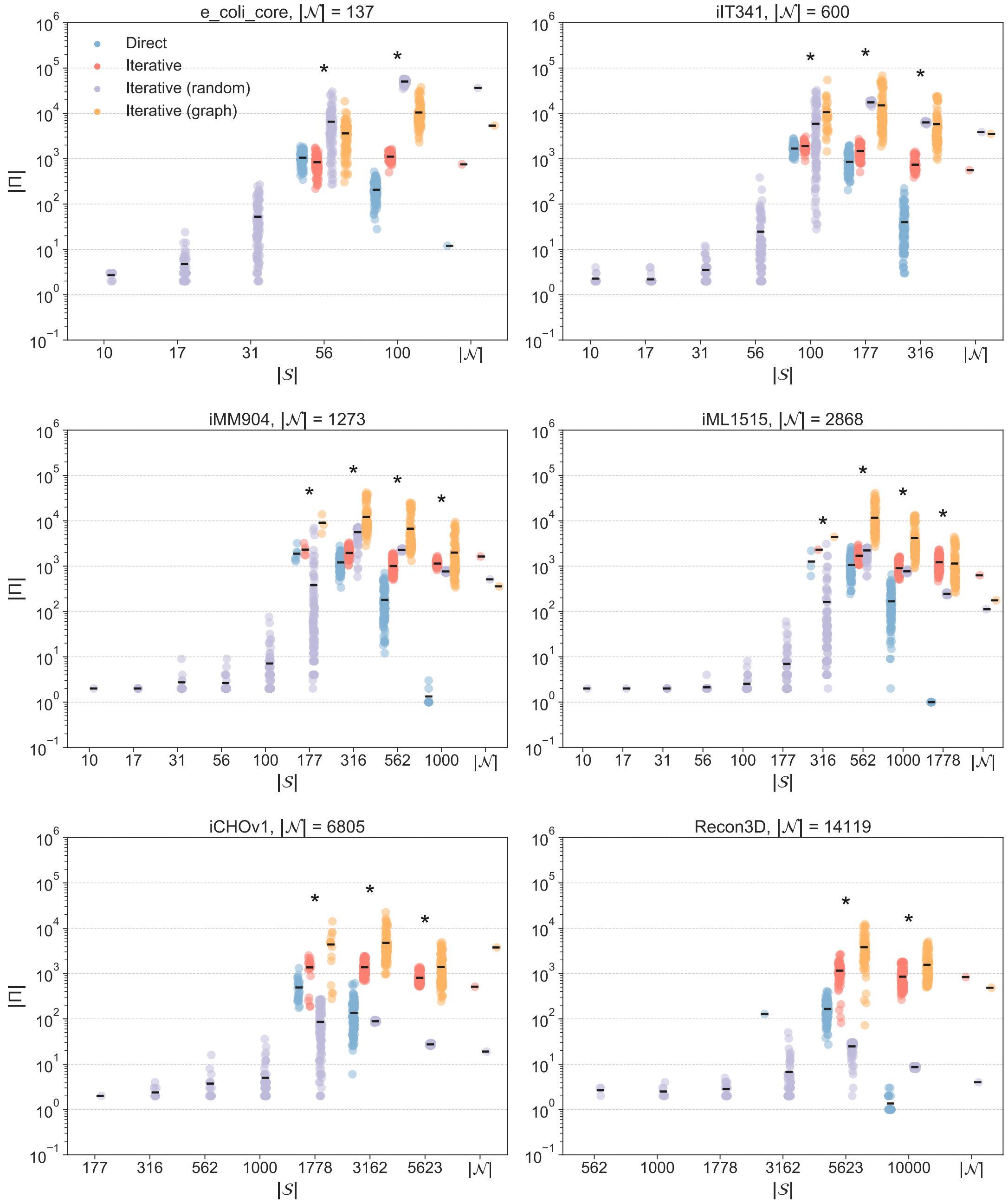


**Figure S3. MPs found in one hour in incomplete enumerations.** Number of MPs |Π| found in one hour with direct minimization (red), iterative minimization (blue), iterative minimization with randomization (purple), and iterative minimization with graph (orange) for different subnetwork sizes |𝒮| in six different models with network size |𝒩|. Stars indicate significant difference between means from one-way ANOVA where data was available for all three methods or *t*-test where data was available for two methods (significance level 0.05). Enumerations that were completed within one hour are not included.

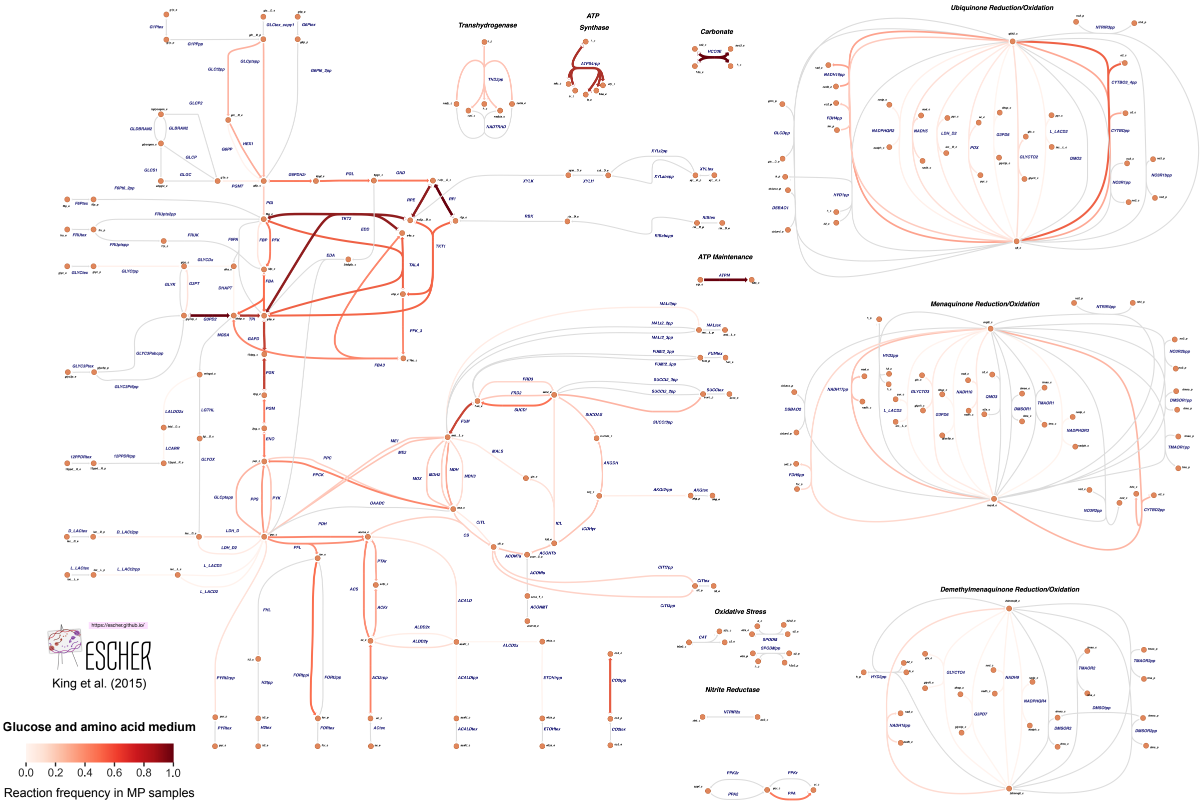


**Figure S4. Pathway map of *E. coli* central carbon metabolism for glucose and amino acid medium.** Circles are metabolites and arrows are reactions. The color and width of arrows indicate reaction frequency across 100,000 randomly sampled MPs. The network was used as a subnetwork within the genome-scale model iJO1366 with aerobic conditions, a glucose and amino acid medium, and growth and ATP maintenance requirements. For reversible reactions, frequency does not distinguish between flux directions. Pathway map made with Escher[54].

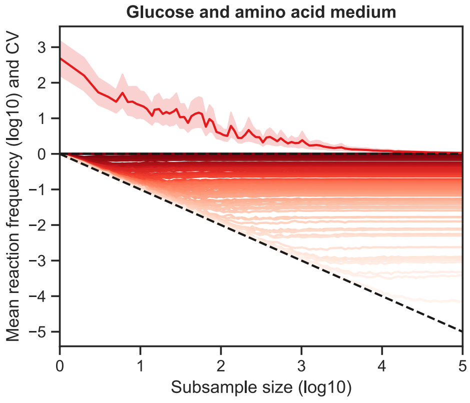


**Figure S5. Convergence of reaction frequencies in MP subsamples for glucose and amino acid medium.** The 100,000 MP samples were subsampled with replacement 100 times for 100 logarithmically spaced subsample sizes ranging from 1 to 100,000. Mean absolute reaction frequency for each reaction is shown below, and mean coefficient of variation (CV) of relative reaction frequency across all reactions is shown above with 95% confidence interval. Dashed lines indicate $y = 0$ and $y = -x$, the line along which reaction frequencies converge with increasing subsample size.

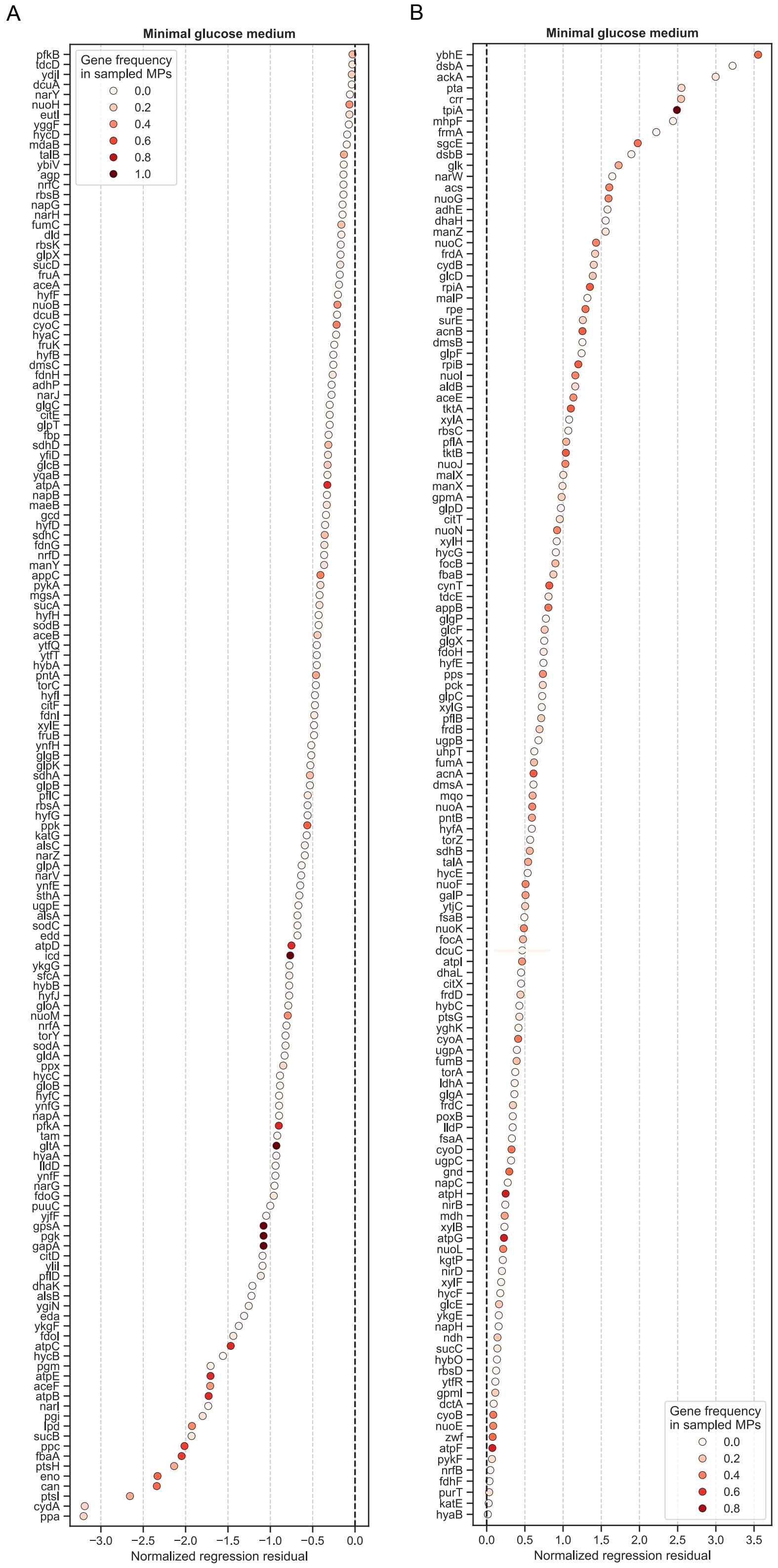


**Figure S6. Regression residuals of single gene knockouts in *E. coli* central carbon metabolism.** (**A**) Normalized residuals of genes with negative residuals (gene frequency in MP samples lower than expected), and (**B**) normalized residuals of genes with positive residuals (gene frequency in MP samples higher than expected), from regression of experimental growth data against gene frequency in MP samples for a minimal glucose medium under aerobic conditions.

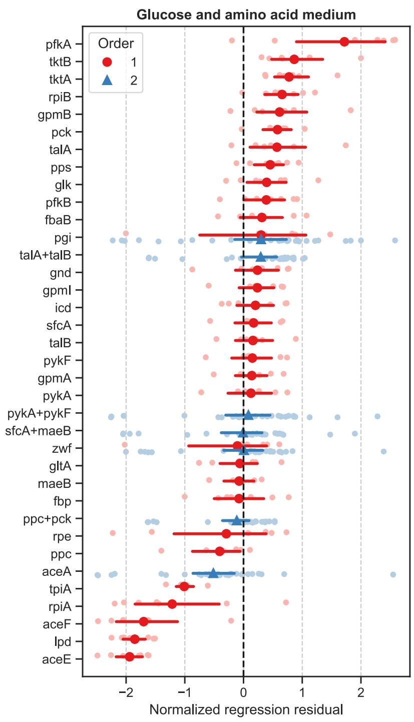


**Figure S7. Regression residuals of multiple gene knockouts in *E. coli* central carbon metabolism.** Normalized residuals of gene sets by order of deletion in multiple gene deletion experiments. Mean is shown for each gene set with 95% confidence interval along with the residuals of all multiple deletions involving the gene set. Some of the secondary deletions include two genes. Negative residuals indicate that gene set frequency in MP samples was lower than expected and vice versa.

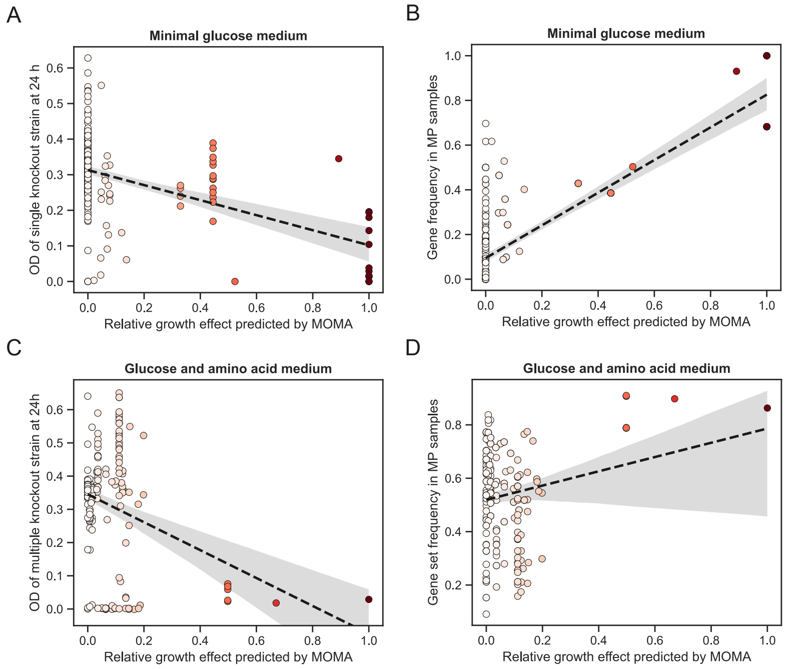


**Figure S8. Correlations of MOMA predictions** Fitted lines with 95% confidence interval from linear regression of relative growth effect predicted by MOMA against (**A**) OD of single gene knockout strains at 24 h and (**B**) gene frequency in MP samples for minimal glucose medium and (**C**) OD of multiple gene knockout strains at 24 h and (**D**) gene frequency in MP samples for glucose and amino acid medium.

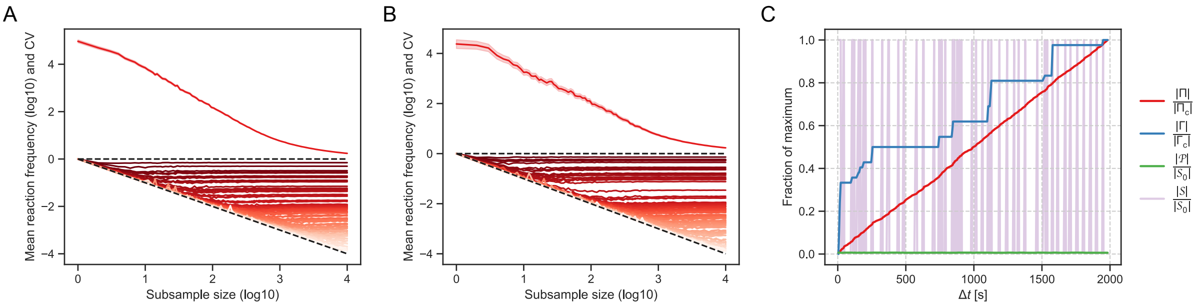


**Figure S9. Convergence of MP sampling and enumeration for sequentially constrained host-microbe model.** Convergence of reaction frequencies in MP subsamples after (**A**) the first step and (**B**) the second step. For each step, the 10,000 MP samples were subsampled with replacement 100 times for 100 logarithmically spaced subsample sizes ranging from 1 to 100,000. Mean absolute reaction frequency for each reaction is shown below, and mean coefficient of variation (CV) of relative reaction frequency across all reactions is shown above with 95% confidence interval. Dashed lines indicate $y = 0$ and $y = -x$, the line along which reaction frequencies converge with increasing subsample size. Trajectories are shown for 100 randomly selected reactions. (**C**) Complete enumeration of MPs and MCSs using iterative minimization with graph after the third step. The number of MPs $|\Pi|$ and MCSs $|\Gamma|$ found is shown along with the MP size $|\mathcal{P}|$ and effective subnetwork size $|\mathcal{S}|$ as a function of running time $\Delta t$. Values are normalized by the size of the complete set of MPs $|\Pi_c|$, the size of the complete set of MCSs $|\Gamma_c|$, or the initial subnetwork size $|\mathcal{S}_0|$.

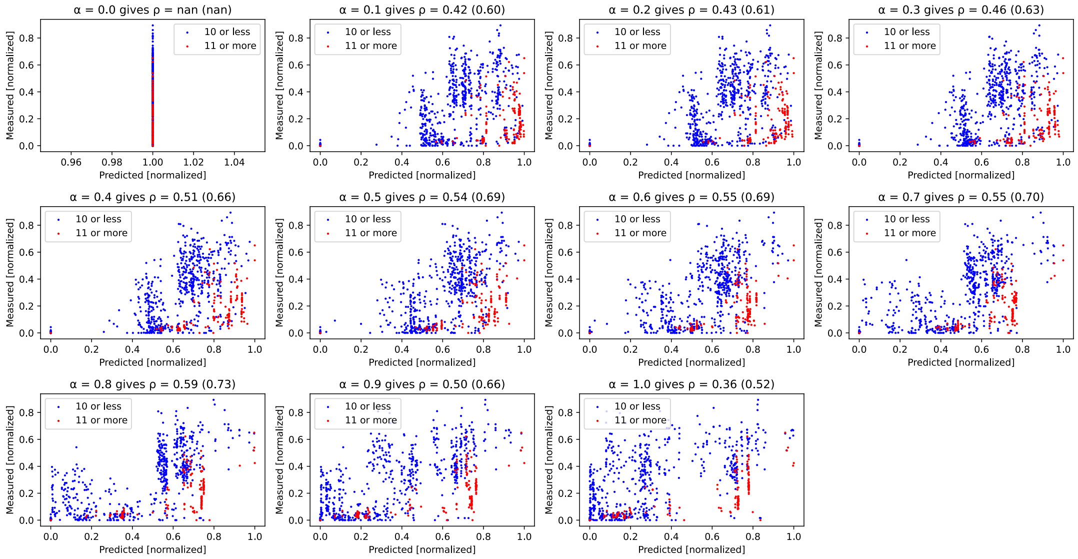


**Figure S10. Simulated versus measured butyrate production** Scatter plots of simulated versus measured butyrate production for 11 linearly spaced values of $\alpha \in [0, 1]$. The simulated and measured values are normalized to $[0, 1]$. Weighted linear correlation $\rho$ between simulations and measurements are given for each plot, with weights equal to the number of times a community was measured. Dot colors display community size. The $\rho$ in brackets denote the correlation obtained when omitting communities with 11 members or more (orange dots).

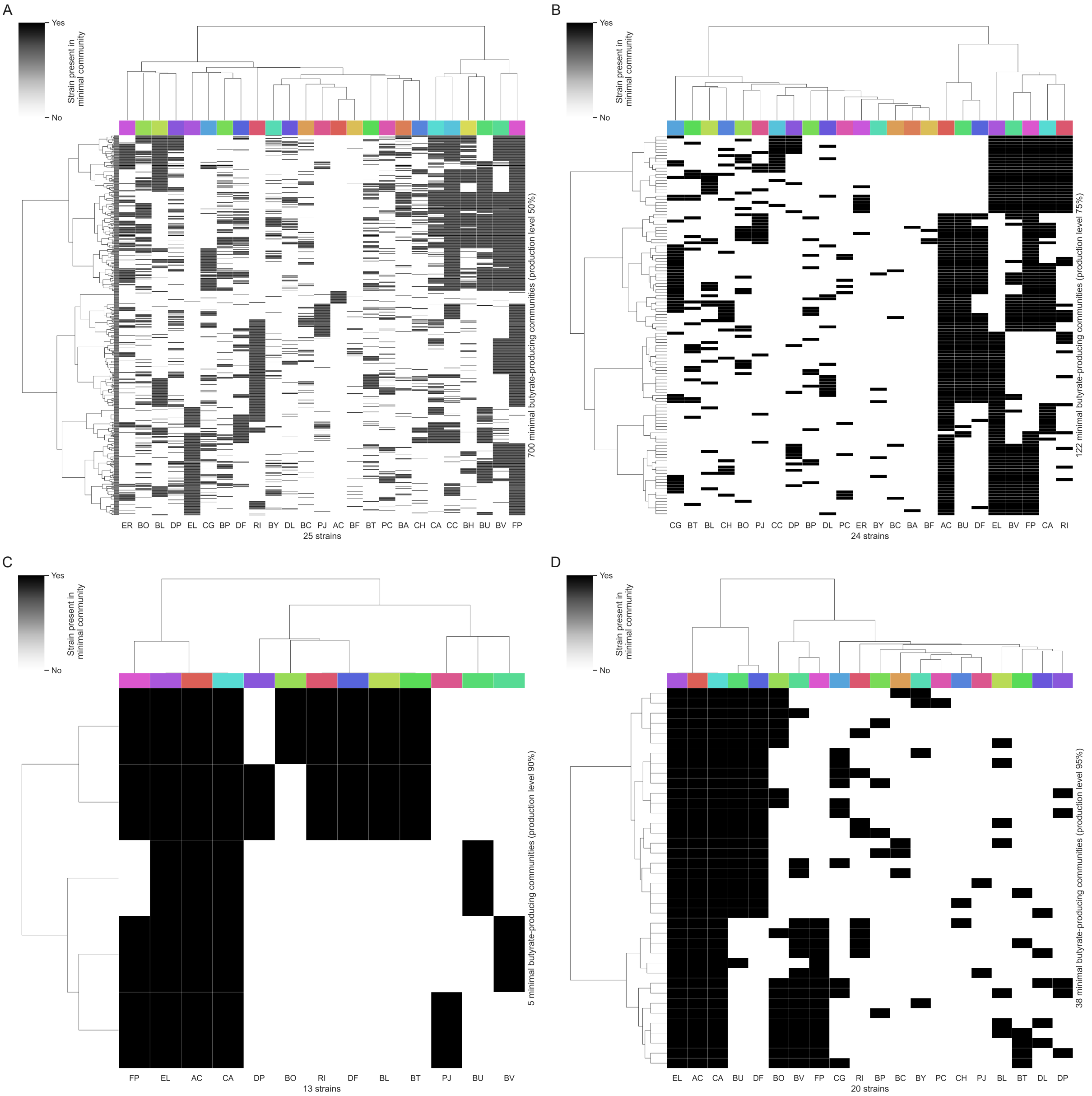


**Figure S11. Minimal butyrate-producing community compositions.** Strain presence in minimal communities enumerated for butyrate production levels (**A**) 50%, (**B**) 75%, (**C**) 90%, and (**D**) 95%. Rows are minimal communities, columns are strains, and each cell indicates whether a strain is present (dark) or absent (light) in a minimal community.

### Supplementary tables

**Table S1.** Models used for benchmarking with number of reactions and metabolites before (and after) preprocessing.

| Organism | Model | Reactions | Metabolites |
|---|---|---|---|
| *Escherichia coli* | e_coli_core | 95 (137) | 72 (72) |
| *Helicobacter pylori* | iIT341 | 554 (600) | 485 (422) |
| *Saccharomyces cerevisiae* | iMM904 | 1,577 (1,273) | 1,226 (722) |
| *Escherichia coli* | iML1515 | 2,712 (2,868) | 1,877 (1,633) |
| *Cricetulus griseus* | iCHOv1 | 6,663 (6,805) | 4,456 (2,749) |
| *Homo sapiens* | Recon3D | 10,600 (14,119) | 5,835 (5,835) |

**Table S2.** Models used in host-microbe model with number of reactions and metabolites.

| ID | Organism | Model | Reactions | Metabolites |
|---|---|---|---|---|
| ST | *Streptococcus thermophilus* | LMG_18311 | 954 | 851 |
| FP | *Faecalibacterium prausnitzii* | A2_165 | 1,225 | 1,047 |
| LL | *Lactococcus lactis* | subsp_lactis_WCFS1 | 1,295 | 1,118 |
| LP | *Lactobacillus plantarum* | Il1403 | 1,753 | 1,348 |
| EC | *Escherichia coli* | str_K_12_substr_MG1655 | 2,313 | 1,625 |
| BT | *Bacteroides thetaiotaomicron* | VPI_5482 | 2,524 | 1,549 |
| HS | *Homo sapiens* | Recon3D | 10,600 | 5,835 |

**Table S3.** Models used in butyrate-producing community enumeration with number of reactions and metabolites.

| ID | Organism | Model | Reactions | Metabolites |
|---|---|---|---|---|
| AC | *Anaerostipes caccae* | DSM_14662 | 1,617 | 1,328 |
| BA | *Bifidobacterium adolescentis* | ATCC_15703 | 1,129 | 991 |
| BC | *Bacteroides caccae* | ATCC_43185 | 2,381 | 1,461 |
| BF | *Bacteroides fragilis* | NCTC_9343 | 2,444 | 1,512 |
| BH | *Blautia hydrogenotrophica* | DSM_10507 | 1,637 | 1,290 |
| BL | *Bifidobacterium longum* | DJO10A | 2,043 | 1,386 |
| BO | *Bacteroides ovatus* | ATCC_8483 | 2,476 | 1,526 |
| BP | *Bifidobacterium pseudocatenulatum* | DSM20438 | 1,583 | 1,185 |
| BT | *Bacteroides thetaiotaomicron* | VPI_5482 | 2,524 | 1,549 |
| BU | *Bacteroides uniformis* | ATCC_8492 | 2,418 | 1,500 |
| BV | *Bacteroides vulgatus* | ATCC_8482 | 2,474 | 1,519 |
| BY | *Bacteroides cellulosilyticus* | DSM_14838 | 2,278 | 1,513 |
| CA | *Collinsella aerofaciens* | ATCC_25986 | 920 | 850 |
| CC | *Coprococcus comes* | ATCC_27758 | 1,599 | 1,280 |
| CG | *Clostridium asparagiforme* | DSM_15981 | 1,960 | 1,462 |
| CH | *Clostridium hiranonis* | DSM_13275 | 1,258 | 1,111 |
| DF | *Dorea formicigenerans* | ATCC_27755 | 2,072 | 1,483 |
| DL | *Dorea longicatena* | DSM_13814 | 1,536 | 1,227 |
| DP | *Desulfovibrio piger* | ATCC_29098 | 980 | 890 |
| EL | *Eggerthella lenta* | DSM_2243 | 1,046 | 961 |
| ER | *Eubacterium rectale* | ATCC_33656 | 1,238 | 1,065 |
| FP | *Faecalibacterium prausnitzii* | A2_165 | 1,225 | 1,047 |
| PC | *Prevotella copri* | DSM_18205 | 1,634 | 1,216 |
| PJ | *Parabacteroides johnsonii* | DSM_18315 | 2,446 | 1,496 |
| RI | *Roseburia intestinalis* | L1_82 | 2,095 | 1,422 |

**Table S4.** Reaction energy of reverse lactose dehydrogenase for two different combinations of reactant concentrations.

| Quantity | Unit | Standard conditions | Realistic conditions |
|---|---|---|---|
| Lactate | mM | $1 \cdot 10^3$ | 1 |
| $NAD^+$ | mM | $1 \cdot 10^3$ | 2.6 |
| NADH | mM | $1 \cdot 10^3$ | 0.083 |
| Pyruvate | mM | $1 \cdot 10^3$ | $2.2 \cdot 10^{-3}$ |
| $\Delta_r G'$ | kJ/mol | $23.7 \pm 0.9$ | $0 \pm 0.9$ |

### Supplementary algorithms

**Algorithm S1:** Finding a new MP with iterative minimization.

**Input:** Subnetwork $\mathcal{S}$ and whether to randomize.
**Output:** New MP $\mathcal{P}$.

```
𝒫 ← ∅;
while 𝒮 ≠ ∅ do
    Deactivate a (random) i ∈ 𝒮;
    𝒮 ← 𝒮 \ {i};
    r ← LP(𝒮);
    if r is optimal then
        if randomize then
            Draw a random j ∈ 𝒮;
            while r_j = 0 do
                Deactivate j;
                𝒮 ← 𝒮 \ {j};
                Draw a random j ∈ 𝒮;
        else
            foreach j ∈ 𝒮 do
                if r_j = 0 then
                    Deactivate j;
                    𝒮 ← 𝒮 \ {j};
    else
        Reactivate i;
        𝒫 ← 𝒫 ∪ {i};
```

**Algorithm S2:** Finding a new viable cut set.

**Input:** Set of known MPS $\Pi$ and set of known MCSs $\Gamma$.
**Output:** New viable cut set $\mathcal{C}$ and set of known MCSs $\Gamma$.

```
repeat
    𝒞 ← BIP(Π, Γ);
    Deactivate all i ∈ 𝒞;
    r ← LP(∅);
    if r is not optimal then
        Reactivate all i ∈ 𝒞;
        Γ ← Γ ∪ {𝒞};
until BIP is infeasible or 𝒞 is not an MCS;
```

**Algorithm S3:** Enumeration of MPs and MCSs with iterative minimization and graph.

**Input:** Subnetwork $\mathcal{S}$.

**Output:** Set of MPs $\Pi$ and set of MCSs $\Gamma$.

$\Pi \leftarrow \Gamma \leftarrow \mathcal{Q} \leftarrow \mathcal{G} \leftarrow \emptyset$;

$\mathcal{S}_0 \leftarrow \mathcal{S}$;

**repeat**

  **if** $\mathcal{Q}$ **then**

    Draw $\mathcal{S}$ from $\mathcal{Q}$;

    $\mathcal{Q} \leftarrow \mathcal{Q} \setminus \{\mathcal{S}\}$;

  **else**

    $\mathcal{S} \leftarrow \mathcal{S}_0$;

  **repeat**

    $\mathcal{C}, \Gamma \leftarrow$ **Algorithm S2**;

    **if** $\mathcal{C}$ *is viable* **then**

      $\mathcal{P} \leftarrow$ **Algorithm S1**;

      $\Pi \leftarrow \Pi \cup \{\mathcal{P}\}$;

      **if** $\mathcal{S} = \mathcal{S}_0$ **then**

        $\mathcal{G} \leftarrow \mathcal{G} \cup \binom{\mathcal{P}}{2}$;

        Find new maximal cliques $\mathcal{K}$ in $\mathcal{G}$;

        $\mathcal{Q} \leftarrow \mathcal{Q} \cup \mathcal{K}$;

  **until** *no more viable cut sets or (*$\mathcal{S} = \mathcal{S}_0$ *and* $\mathcal{Q}$*)*;

**until** $\mathcal{S} = \mathcal{S}_0$ *and BIP is infeasible*;
